# Orthogonality of sensory and contextual categorical dynamics embedded in a continuum of responses from the second somatosensory cortex

**DOI:** 10.1101/2023.09.22.559070

**Authors:** Lucas Bayones, Antonio Zainos, Manuel Alvarez, Ranulfo Romo, Alessio Franci, Román Rossi-Pool

**Affiliations:** Instituto de Fisiología Celular ─ Neurociencias, Universidad Nacional Autónoma de México, 04510 Mexico City, Mexico; El Colegio Nacional, 06020 Mexico City, Mexico; Departmento de Matemática, Facultad de Ciencias, Universidad Nacional Autónoma de México, 04510 Mexico City, Mexico; Montefiore Institute, University of Liège, Av de la Découverte 10, 4000 Liège, Belgique; WEL Research Institute, avenue Pasteur, 6, 1300 Wavre, Belgique; Centro de Ciencias de la Complejidad, Universidad Nacional Autónoma de México, Mexico City, Mexico

## Abstract

How does the brain simultaneously process signals that bring complementary information, like raw sensory signals and their transformed counterparts, without any disruptive interference? Contemporary research underscores the brain’
ss adeptness in using decorrelated responses to reduce such interference. Both neurophysiological findings and artificial neural networks (ANNs) support the notion of orthogonal representation for signal differentiation and parallel processing. Yet, where, and how raw sensory signals are transformed into more abstract representations remains unclear. Using a temporal pattern discrimination task (TPDT) in trained monkeys, we revealed that the second somatosensory cortex (S2) efficiently segregates faithful and transformed neural responses into orthogonal subspaces. Importantly, S2 population encoding for transformed signals, but not for faithful ones, disappeared during a non-demanding version of the task, which suggests that signal transformation and their decoding from downstream areas are only active on-demand. A mechanistic computation model points to gain modulation as a possible biological mechanism for the observed context-dependent computation. Furthermore, individual neural activities that underlie the orthogonal population representations exhibited a continuum of responses, with no well-determined clusters. These findings advocate that the brain, while employing a continuum of heterogeneous neural responses, splits population signals into orthogonal subspaces in a context-dependent fashion to enhance robustness, performance, and improve coding efficiency.

**SIGNIFICANCE STATEMENT:** An important function of the brain is turning sensation into perception. Yet, how this function is implemented remains unknown. Current research, insights from artificial neural networks, highlights using of orthogonal representations as an effective means to transform sensory signals into perceptual signals while separating and simultaneously processing the two information streams. Neuronal recordings in S2 while trained monkeys performed the TPDT, revealed that this function is implemented at the population level. While S2 encodes sensory information independently of context, the encoding of categorical information, like task parameters, is only performed when the task demands it. Such distinct and flexible organization, enriched by a spectrum of neural activities, reflects the brain’s efficiency, resilience, and overall purpose for solving cognitive tasks.

## Introduction

The kitchen, a commonplace yet remarkably complex sensorial arena, serves as a vivid illustration of the extraordinary capabilities of the human brain. From the moment one steps into this space with the intent to prepare a meal, a plethora of sensory information floods our neural pathways - the visual recognition of ingredients and utensils, the auditory distinction between background hums and salient sizzles, the tactile feedback of knife against cutting board, the olfactory delight of sautéing garlic, and the gustatory assessment of flavor. Within this sensory tapestry, the brain performs a series of intricate transformations, transducing raw stimuli into meaningful units of information, and translating them into coherent, goal-oriented actions. This process encapsulates the remarkable ability of our neural networks to process diverse sensory information and seamlessly integrate them into a single, coherent experience, an ability that lies at the core of human cognition and behavior. Nevertheless, this raises an intriguing question: how does the brain simultaneously receive raw sensory signals, process them into more abstract transformed ones, and achieve representation and inter-area transmission of both kinds of signals in a context-dependent fashion?

Experiments on behaving monkeys has brought forth an attractive hypothesis about the brain’s unique encoding mechanisms (1–5). It suggests that the brain utilizes orthogonal encoding subspaces, a kind of cognitive architecture, to reduce interference among signals involved in different processes. It has been seen that this smart arrangement allowed the brain to decouple mnemonic coding with dynamics related to stimulus representation and comparison (4, 5). Furthermore, it also permitted the network to separate signals associated with motor planning and motor execution (1–3). Moreover, orthogonal subspaces were also found in tasks involving sequential memory representation (6) and between signals that encode reward and confidence (7). Further research in monkeys has revealed that the neocortical neuronal network can represent multiple related task variables without cross-interference (6). Additionally, studies conducted in the posterior parietal cortex of rodents, demonstrated that the activity patterns of neuronal groups during the evidence accumulation process are significantly influenced by past events (8). Here, the neuronal network employs a degree of orthogonalization to distinguish and reduce interference between learned information and new experiences. Complementarily, recent work in rodents suggested that in order to reduce correlation in the neural representations of related task variables, the population must code the different parameters into decoupled dynamics (9). Therefore, the notion of orthogonal representation has garnered support from a variety of studies. However, it remains unclear whether this type of neural scaffold is effective in distinguishing between raw sensory signals, originating from stimulus input, and transformed signals, which represent sensory information that has been processed and abstracted for use by downstream areas in the hierarchical processing pathway.

Research in artificial intelligence, particularly with artificial neural networks (ANN), has substantially deepened our understanding of orthogonal representations in signal discrimination. Much like their biological counterparts, ANNs have been observed to use orthogonal representations for context-specific tasks (10). Recent studies in spiking neuronal networks indicate that leveraging latent orthogonal subspaces can boost learning versatility (11). ANNs have also adeptly mimicked outcomes from biological data across several tasks. This demonstrates their capability to distinctly showcase orthogonal differentiation in input, scaling, and gain subspaces (12–14), as well as in category dynamics during delay periods (15). A notable contemporary finding is that both biological and artificial networks encode neural noise and stimulus coding in orthogonal subspaces. This maximizes the decorrelation between population dynamics, thereby enhancing the precision of relevant coding (16). In essence, the employment of artificial networks has harnessed the power of orthogonal representations, revealing promising avenues for enhanced signal processing and task-specific performance.

While the concept of orthogonal encoding and its role in cognitive processes have been extensively studied, little attention has been given to investigating how faithful sensory signals interact with transformed dynamics. Among the variety of brain areas, the secondary somatosensory cortex (S2) constitutes a promising circuit to examine this issue. In our prior research (17), involving macaques trained to undertake a temporal pattern discrimination task (TPDT), we revealed that S2 showcases a broad spectrum of neuronal responses. These ranged from faithful or phase-locked responses to more abstract representations, wherein sensory information from stimuli was integrated and transformed by neurons into direct category encoding. Employing information theory at the single-cell level, we noted that these responses span a continuum, moving from sensory to categorical encoding. Building on this work of single-cell analysis, in this study we delve into how these signals might interact at population level. This investigation could yield critical insights about the mechanisms underlying sensory representation and its transformation, aiming to minimize interference among dynamics at the network level. Our exploration of S2’s dynamics will not only contribute to a broader knowledge of how the brain converts sensory inputs but also shed light on the context dependent recruitment of decorrelated population signals during cognitive tasks. This research offers insights that could potentially overcome the constraints of previous studies that used single units in S2 (17–19). By using the same data collected through the TPDT, in which monkeys had to compare if the temporal pattern of two stimuli sequentially presented were equal or different, we found that faithful and categorical dynamics appeared decoupled in the S2 network. Hence, decorrelated combinations of neural responses are necessary to decode sensory inputs and categorical and more abstract signals. Moreover, by employing a non-linear dimensionality reduction method [Uniform Manifold Approximation Projection (UMAP)], renowned for its ability to discern continuity or clusterization of high-dimensional data structures, we showed that the population dynamics elicited by the S2 network arise from a continuum of responses, rather than isolated clusters of neurons. The observed continuity starkly differed in a less-demanding version of the task (the light control variant), where the categorically distinct population signal faded, uncovering a discontinuity in the underlying response patterns. This indicates that such responses manifest exclusively under necessary conditions. To provide mechanistic insights, we present a computational model that serves as an example of the possible dynamical mechanisms underlying this phenomenon in a way that is compatible with the orthogonalization of coding subspaces found in the S2 network.

## Results

### Individual neuronal responses in S2 during task performance

Two monkeys were trained in the TPDT (20), in which they had to report whether two temporal patterns with a vibrotactile flutter stimuli architecture (P1 and P2), were the same (P1=P2) or different (P1≠P2) (Fig. 1A, see Methods). Two possible temporal patterns were presented: extended (E), made of 5 pulses uniformly spaced in time, and grouped (G), with 3 pulses grouped at the middle of the stimulus. There were therefore 4 classes of possible temporal pattern pairs: G-G (c1, red color), G-E (c2, orange), E-G (c3, green) and E-E (c4, blue). Mean stimulus frequency (5 Hz) and duration (1 s) were maintained constant throughout the task. The average performance in the TPDT across S2 recording sessions was 84% (± 7%), and this was also consistent across classes (Fig. 1B).

**Figure 1.**
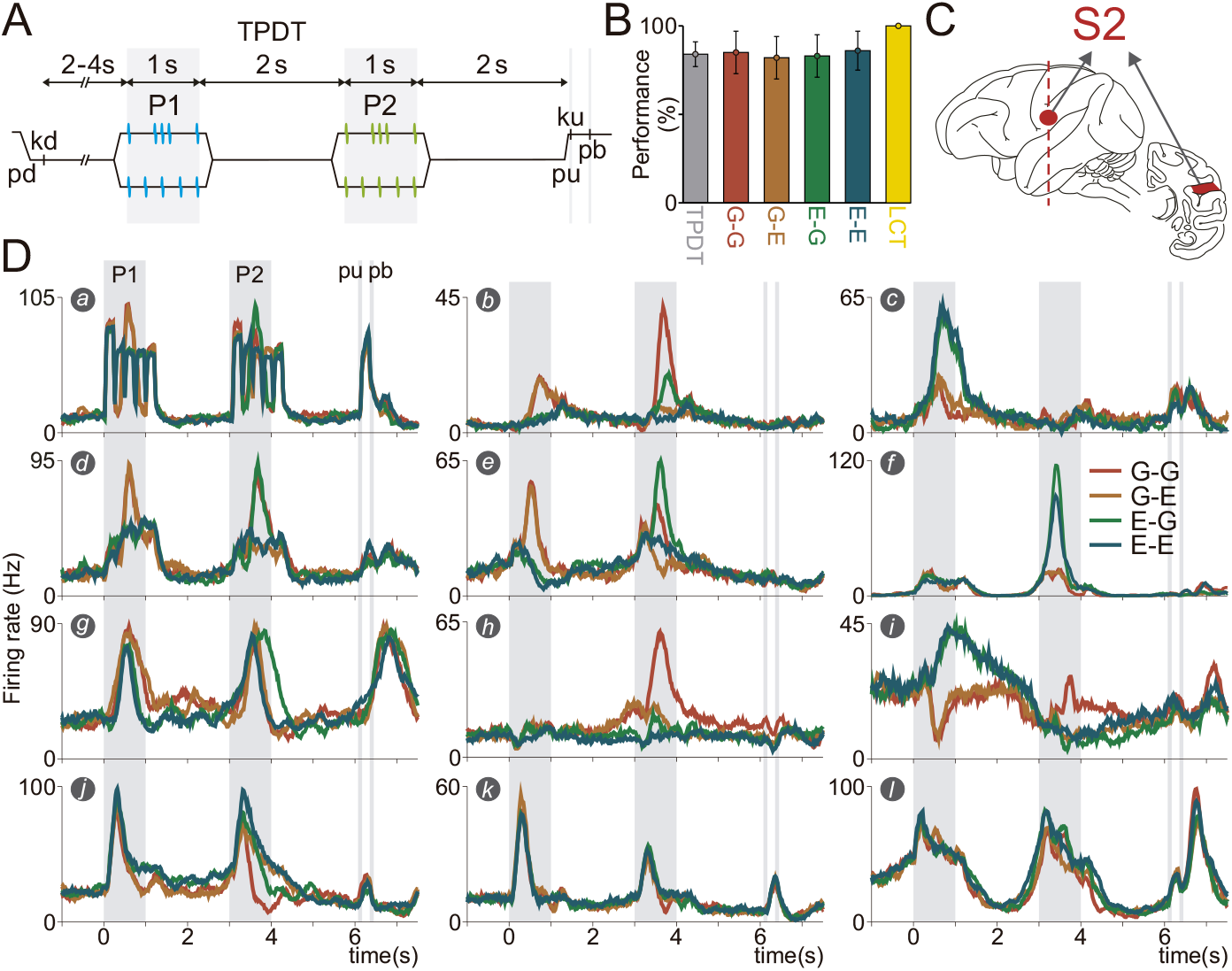
Task, performance and exemplary responses of S2 neurons. (A) Schematic of the temporal pattern discrimination task (TPDT) with trial event sequence. A mechanical probe was lowered (pd), indenting the glabrous skin of one fingertip of the monkey’
ss restrained right hand. Consequently, the animal responded by placing its left free hand on a fixed key (kd). After a variable prestimulus period (2 to 4 s), the probe executed the first vibrotactile stimulus (P1, 1 s duration and mean frequency of 5 Hz) with two possible patterns: grouped (G) or extended (E). After a first fixed delay of 2 s (from second 1 to 3), the second stimulus (P2) was presented, with the same configuration of P1 (1 s duration and G or E patterns). Then, a second fixed delay (2 s) was presented (from second 4 to 6), which ended with the probe up event (pu). After “pu”, the monkey released its free left hand from the key (ku) and pressed one of two buttons (pb) to report its decision, lateral to indicate that P1 and P2 were the same (P1=P2) and medial to indicate that P1 and P2 were different (P1≠P2). (B) Performance for the whole TPDT (gray, n= 423 sessions), for each class (G-G (red), G-E (orange), E-G (green) and E-E (blue)) and the whole light control task (LCT) (n= 76 sessions). (C) Illustration of the brain left hemisphere (left figurine) and a coronal brain slice (right figurine) with S2 (recorded area) highlighted in red. Recordings were made contralateral and ipsilateral to the stimulated fingertip (D) Average firing rates of twelve exemplary neurons, separated per class, plotted in the whole TPDT. Color traces indicate the four possible classes: G-G (red), G-E (orange), E-G (green) and E-E (blue). Note that unit *a* displays purely sensory dynamics whereas units *c, h & f* show highly categorical response. Some neurons displaying mixed signals (sensory and categorical) and temporal responses, can also be seen in units *d, e* and *k, l*, respectively.

We recorded the activity from 1646 neurons in S2 (Fig. 1C, see Methods) during the TPDT (Monkey RR17, n=1035; Monkey RR20, n=611). The responses of 12 and 24 exemplary S2 neurons are displayed in Fig. 1D and Supplementary Fig. 1, respectively. As we previously showed (17), neurons from this area exhibit a broad repertoire of responses with distinguishable neuronal dynamics. Several neurons responded only when sensory stimulation was present and their response faithfully represented the stimulus patterns (some examples in Fig. 1D is *a*, and in Fig. S1 are *a & b*). These neurons were therefore cataloged as “sensory”, “faithful” or “phase-locked”. Other recorded neurons, cataloged as “categorical”, exhibited a non-sensory kind of response. In those neurons, sensory information was transformed to encode stimuli categories or some other features of the task, rather than the faithful temporal course of stimulus. If we look at these categorical units, they tend to respond more strongly to specific stimulus patterns (G or E) or particular classes (some examples in Fig. 1D are *c, e, f & h*, and in Fig. S1 are *I, j, p & q*). For example, unit *b* in Fig. 1D responded more strongly when the G pattern was presented during P1, whereas during P2 the same neuron responded selectively to classes c1 & c3. Furthermore, looking at unit *f* in Fig. 1D, one can notice that during P2 this neuron responded preferentially to classes c3 & c4 rather than c1 & c2. Moreover, some neurons exhibited a mixture of sensory and categorical responses and were cataloged as “mixed”. For instance, neuron *c* in Fig. S1, faithfully represented all classes during P1 but selectively diminished its activity in response to class c1 during P2. More neurons exhibiting a mixture of sensory and categorical responses are unit *d, g & j* in Fig. 1D and units *m, f*, and *s* in Fig. S1. If a sensory or categorical neuron firing rate increased (or decreased) in response to stimulation, it was labeled as positive (or negative, Fig. S1 *k & u* and Fig. S2 *c*). It has been previously reported that the integration of positive and negative S2 neuronal pools improve stimulus information decoding (21, 22). Finally, a fourth set of neurons exhibited strong temporal dynamics (Fig. 1D *k & l*, and Fig. S1 *v, w & x*). These units behaved like a timekeeper, responding independently of the stimuli. It has been speculated that these temporal responses constitute an essential underlying process for the network to predict the arrival of relevant sensory events (23). In the light of this heterogeneous collection of neuronal activity, one thing becomes clear: S2 exhibits a broad range of neuronal signals, representing sensory signals in disparate ways, from purely sensory to purely categorical and in between. This observation makes S2 a natural candidate for a place where sensory and categorical representations encounter, and perception begins.

All the neurons recorded in the light control task (LCT, n=313, of which n=189 from Monkey RR17 and n=124 from Monkey RR20) were also recorded in the TPDT. This pool of neurons, recorded in both experimental conditions, provides a direct comparison of how cognitive demand (high in TPDT and low in LCT) modulates neuronal activity at the single cell level (17). In each trial, monkeys received the same stimuli as in the TPDT, but the correct choice was indicated by a visual cue (see “Methods”). The animal’s performance was 100% in the control variant (Fig. 1B, yellow bar), indicating a strongly reduced cognitive demand. Supplementary Figure 2 displays 16 neurons recorded both in TPDT and LCT. It is possible to observe that purely sensory units were not affected by the change of context (Fig. S2 *a & c*), in line with what we recently reported (17). On the other hand, the response of purely categorical units in TPDT became completely unresponsive to task parameters in LCT (Fig.S2, *g, k, n & p*). This is also in agreement with our analysis of single unit activity in the dorsal premotor cortex (DPC), where categorical units silenced significatively their responses during LCT (20). Finally, mixed units only partially altered their coding during LCT. Specifically, they tended to retain the sensory component while losing the categorical one. For example, unit *e* in Fig.S2 responded selectively to the G pattern during P2 in the TPDT but completely lost its categorical representation in LCT, leaving only the sensory trace of the neuron’s response. Summarizing, S2 neurons can modify or silence their categorical representation in a context-dependent manner, independently of whether they also present a sensory response. This supports the hypothesis that categorical coding is predominantly activated in contexts with high cognitive requirements (TPDT), highlighting its role in information processing that requires abstract interpretation and decision-making based on perceived stimuli. Conversely, sensory coding, which directly reflects the physical properties of stimuli, is active independently of the cognitive demands required by the context. This suggests that sensory coding operates independently of task complexity, providing a stable foundation for perception regardless of the context.

### Cognitive demanding dynamics represented in S2

To evaluate response variability associated with task parameter coding we computed the population instantaneous coding variance during TPDT and LCT (*Var Cod*, Fig. 2 A & B, blue trace). This metric evaluates the variability in neural response rates among different classes and individual neurons at each time point. A high variance in coding (*Var Cod*) indicates that distinct task parameters elicit significantly distinct neuronal activities. We examined the response variance elicited by different task parameters, namely, the first stimulus identity (*Var P1*), the second stimulus identity (*Var P2*), and the decision (*Var Dec*) (Fig. 2 A & B, light blue, purple and pink, traces, respectively). *Var P1* therefore measures the response variability emerging during the first stimulus period, *Var P2* the response variability during the second stimulus or comparison period, and *Var Dec* the response variability emerging during the decision-making period. The variances in the peristimulus period reflect the basal fluctuations across trials (∼ 2 sp/s). Parameter variances higher than their respective basal values are associated with relevant coding. To facilitate the comparison across parameters, values associated to basal variance were subtracted in each kind of variance computed. In the TPDT, *Var Cod* and *Var P1* traces showed significant coding for sensory representation during P1. Importantly, the *Var P1* wax and wane during the delay period, exhibiting a recall of P1 information during the comparison period. In addition to *Var P1, Var P2* and *Var Dec* increase significantly during P2. As opposed to what we observed in DPC (23), where only a small percentage of neurons encoded the second stimulus, S2’
ss neurons showed a marked coding of the identity of P2. The sharp increment in *Var Cod* during P2, is caused by the appearance of specific class and decision coding. Particularly, as we have shown previously, several neurons respond differently for a particular class during P2 (see neurons *n* and *p*, in Fig. S2). Finally, immediately after “pb”, significant decision-related variance emerged (*Var Dec*). Notably, all these cognitive-dependent fluctuations vanished in the LCT (Fig. 2B). Given that the correct report is indicated by a persistent visual cue, only variances related to precise stimulus identity representation remained in LCT, depicted by increments of *Var P1 & Var P2* in its corresponding periods. Further, *Var P1* dynamic across the delay and P2 vanished; *Var P2* was reduced noticeably; and *Var Dec* faded away. These results suggest that while phase locking responses remained during the control condition, categorical and decision coding disappeared (17). This evidence reveals once again the true nature of S2, a colorful palette of sensory and cognitive demanding categorical dynamics, with a striking mixture in-between.

**Figure 2.**
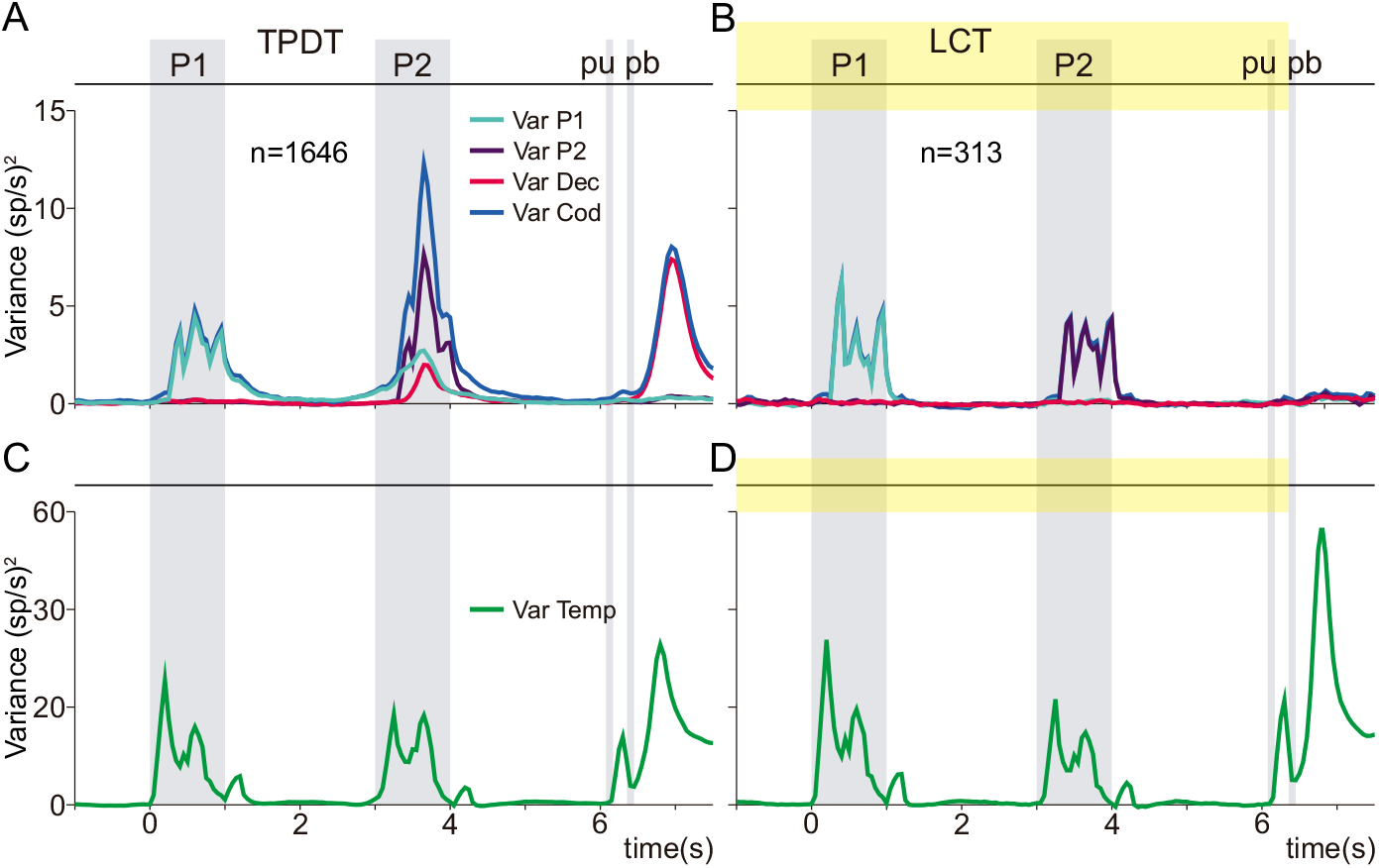
Population variance during active and control tasks. (A, B) Population variances as a function of time during the TPDT (n=1646) and LCT (n=313). Traces refer to P1 (*Var P1*, light blue trace), P2 (*Var P2*, purple trace), decision (*Var Dec*, pink trace) and coding (*Var Cod*, blue trace) variances. Note that basal variances (associated with residual fluctuations) were subtracted in all variances computed. Akin to frontal lobe areas, categorical and perceptual responses are abolished when LCT is employed, but sensory representations related with stimulation persist. (C, D) Temporal (or non-coding) variance during TPDT and LCT (*Var Temp*, green trace). This metric captured the sequential events of the task rather than across classes. Note how under LCT the temporal variance increases with respect to TPDT, meaning that non-coding signals constitute an essential part of the network dynamics.

Single unit activities reported in Figs. 1 and S1 revealed that a significant number of neurons exhibited a purely temporal, i.e., independent of task parameters, behavior. To investigate the amount of population variance associated with this kind of dynamics during TPDT and LCT, we calculated the instantaneous variance with respect to the temporal-averaged response (*Var Temp*, Fig. 2C and D, green trace). Whenever the population activity deviates from its average value across the task period the temporal variance will increase. Interestingly, during TPDT, the instantaneous temporal population variance was much higher than the cognitive variances at any time bin. This indicates that most of the population variance over the course of a trial captured the variability across sequential events of the task rather than across classes. Additionally, under the LCT condition, the instantaneous temporal variance is even higher in comparison to TPDT. This means that non-coding (or temporal) signals constitute an essential part of the network dynamics.

To further explore the nature of the coding signals, we categorized the recorded neurons based on their location in either the left (n=1162) or right (n=484) brain hemispheres and then calculated the same population variance measures as Fig. 2 (Fig. S4 A & C). Population variance measures were similar in the two hemispheres. There were however small differences as well. The left hemisphere (ipsilateral to stimulation) exhibited more pronounced sensory-shaped dynamics compared to the right hemisphere. In contrast, the right hemisphere displayed a notable increase in *Var Dec* relative to the left, which aligns with expectations given its ipsilateral position to the motor execution. The evaluation of condition-averaged dynamics (*Var Temp*) revealed no substantial differences between hemispheres, except for a minor increase in this metric during stimulus periods for the left hemisphere, and a more consistent activity after the “pb” event in the right hemisphere.

This evidence further underscores the multifaceted essence of S2, portraying it as a vibrant mosaic of sensory inputs and cognitively demanding categorical dynamics. The findings highlight a striking interplay between these elements, accentuated by the nuanced differences in hemisphere involvement, which adds depth to our understanding of S2’s role in integrating and processing complex sensory and decision-related information.

### Orthogonal sensory and categorical dynamics in S2

To further explore the interaction between different types of coding across the heterogeneous responses of the neuronal population of S2, we applied dimensionality reduction methods to unravel the latent dynamics behind these responses. The S2 population response can be represented as a point evolving in time in a *n*-dimensional Euclidean space, where *n* is the number of recorded neurons (TPDT, n=1646; LCT, n=313). We used the classical meta-population approach, in which the firing rate of separately recorded neurons are combined into a population vector of firing rates (4–7, 24, 25). As neuronal activity evolves over time, the point moves through the *n*-dimensional space, creating a trajectory that represents the population response. Principal component analysis (PCA) can be used to identify the linear low-dimensional subspace capturing the most salient features (principal components, PCs) of the population dynamic response. However, as we have pointed out, much of this population variance is related to temporal dynamics. Therefore, to study the different types of relevant coding across the population, it might be useful to find a way to diminish the impact of those non-coding signals. For this, one possible solution is to apply PCA to the population responses but limiting it to a narrow period of the task (5, 26). In this case, the temporal variance is much smaller, and the covariance matrix is determined by the coding correlation across neurons throughout this period. Therefore, we applied PCA to the population covariance matrices computed during the final 0.6 seconds of the stimulus periods: P1 and P2, as well as the last 1.5 seconds of the task, which corresponded to the push button period (Fig. 3A, B & C, pink marker on top). Again, by focusing on a small period of the task, we can effectively reduce temporal variance and isolate key aspects of the dynamics. Subsequently, we projected neural activity during TPDT and LCT onto these PCs that were ordered according to their explained total variance.

**Figure 3.**
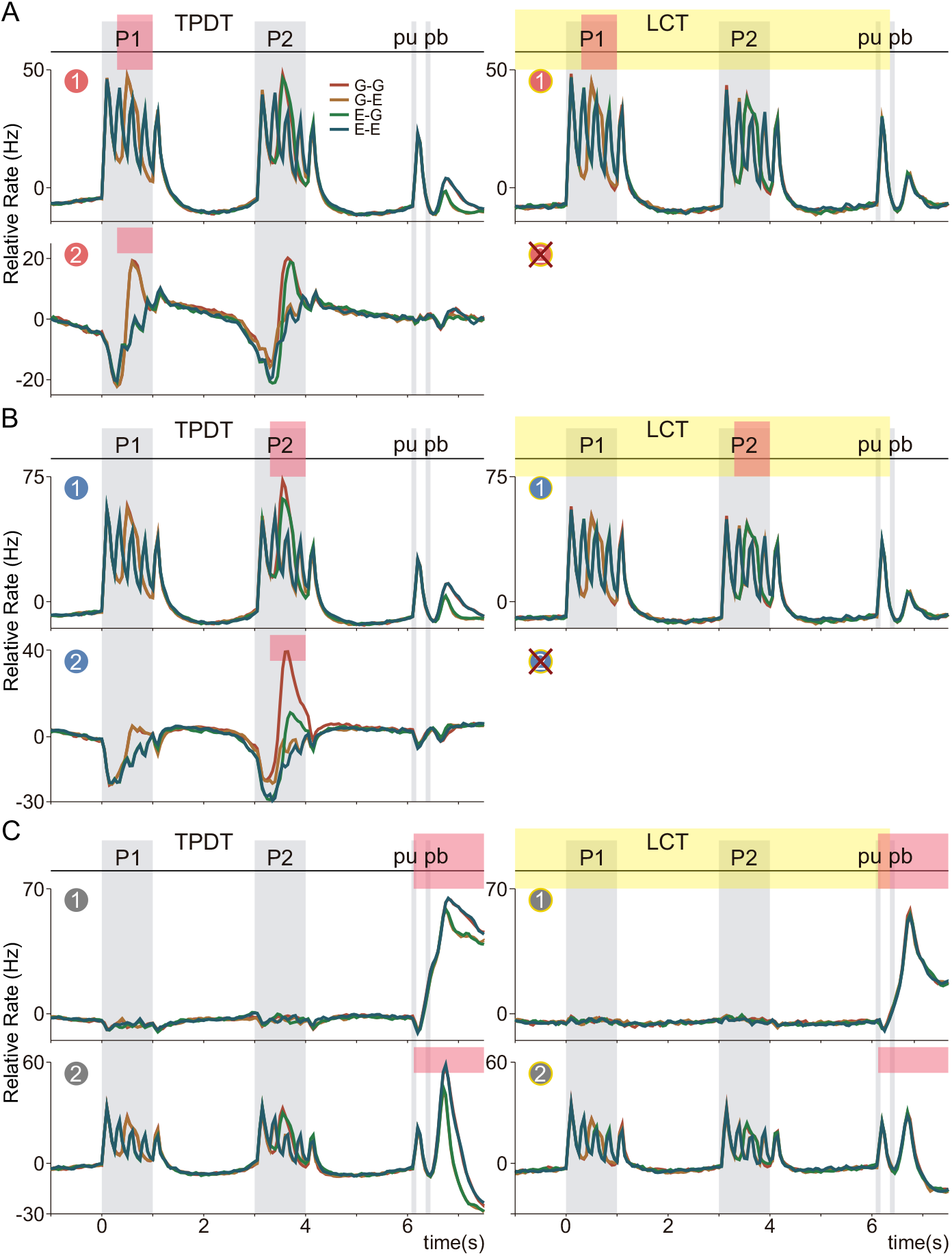
Contextual orthogonality in population dynamics. (A, B and C) Principal component analysis (PCA) was applied to the covariance matrices computed on neural activity during the last 0.6 s of P1 and P2, and 1.5 s of the push button period (pink markers on top), to compute associated principal components (PCs) P1-PCs, P2-PCs and pb-PCs. The activity from the whole (−1 to 7.5 s) TPDT (n=1646) and LCT (n= 313), sorted by class identity, was projected onto the resulting PC axes and ordered by their explained variance (ETV). In TPDT only the first two PCs for each parameter (1 & 2 encircled numbers) were above the noise level, while in LCT this condition was met only for P1-PC1, P2-PC1 and the first two dec-PCs. (A, B) TPDT projections with P1-PC1 and P2-PC1 exhibited strong sensory dynamics, which were barely unchanged in the LCT, whereas dynamics obtained with P1-PC2 and P2-PC2 displayed more transformed and categorical-related responses. The latter ones completely disappeared in the LCT. (C) TPDT projections onto pb-PC1 and PC2 showed dynamics purely related with decision during the intertrial period. In LCT only the temporal component of this dynamics remained.

This period-restricted PCA analysis returned three sets of PCs, respectively associated with P1 (P1-PCs), P2 (P2-PCs) and push-button (pb-PCs). Only the significant PCs that explained more variance than a permuted covariance matrix are shown in Fig. 3 (see Methods). Notice that in the control variant only one PC is significant for both stimulus periods. When we projected the whole TPDT (−1 to 7.5s) on P1-PCs and P2-PCs, we noticed that the first PCs (Fig. 3A & B, P1-PC1 & P2-PC1, respectively) captured strong sensory (phase-locked) signals. Conversely, the second principal components in both cases (Fig. 3A & B, P1-PC2 & P2-PC2, respectively) encapsulated responses that do not faithfully represent sensory stimuli but rather discern some of their categorical properties. In particular, P1-PC2 discerned the extended or grouped identity of both the first and second stimulus, whereas P2-PC2 mostly discerned the class of the second stimulus by exhibiting a large response only in the G-G class. It is crucial to emphasize the dynamics driven by P2-PC2, which are closely associated with class coding (categorical coding), particularly during the P2 period (Fig. 3B). This pattern aligns with the increase in *Var Cod* observed in Fig. 2 A, suggesting that these dynamics likely emerge for the purposes of comparison. Remarkably, despite the axis having been computed with an unsupervised approach, sensory and categorical signals manifested in different components. This means that the population of S2 neurons intrinsically represent the faithful and categorical responses as orthogonal dynamics. Notice, in particular, that during LCT categorical responses disappeared and only the faithful ones remained (right panels in Fig. 3A & B).

To further corroborate this separation between sensory and categorical dynamics, we applied PCA to the covariance matrix quantified from the whole (−1s to 7.5s) TPDT and LCT (Fig. S3 A & B). At first glance, PCA revealed a complex picture of population dynamics, with both sensory and categorical signals distinctly emerging. Akin to those observed in Fig. 3, the first PC demonstrated a faithful sensory response, while the third PCs exhibited categorical dynamics (Fig.S3 A). This result supports the hypothesis that sensory and categorical signals are decoupled at the population level. In agreement with previous research, our results suggest that orthogonality in population dynamics might be employed by the S2 network during the TPDT (3, 4, 6). The resulting decoupled dynamics might act as an efficient mechanism to transform sensory inputs into categories while protecting from sensory interference. Further, when PCA was applied to the LCT (Fig.S3 B), only the sensory component and the dynamic during PB were significant with respect to noise. Analogously to what we observed in single units, PCs in LCT displayed either purely sensory (PC1) or temporal responses (PC2). Thus, the orthogonal categorical signals arise only when the task is cognitively demanding. Finally, when focusing on the last period of the task, pb-PCs (Fig. 3C, pb-PC1 & pb-PC2) displayed responses akin to PC2 & PC4 in Fig. S3. These components exhibit a pronounced decision-related response, delineated by categorical (matched/non-matched classes) distinctions following the pb event, which is in line with the significant increase in *Var Dec* observed in Fig. 2A. In the LCT, only the temporal component of the decision remained during the push-button period (Fig. 3C & Fig. S3B), which confirms the reduced cognitive effort in the LCT.

The organization of sensory, categorical, and decision-related dynamics into orthogonal subspaces, as depicted in Fig. 3 and S3, is similarly maintained across brain hemispheres at the population level (Fig. S4 B & D). In both hemispheres, the 1st and 3rd PCs predominantly capture sensory and categorical dynamics, respectively, whereas the second PC reflects a combination of decision-related responses with a minor sensory component. Although no significant regional differences were observed in the types of emergent signals, the left hemisphere showed a modest enhancement in the intensity of both sensory and categorical dynamics. Conversely, the right hemisphere demonstrated more pronounced decision-related dynamics following the “pb” event. This observation aligns with the variance analysis in Fig. S4 A & C, revealing a coherent picture across different levels of analysis.

Overall, the PCs resulting from the period-restricted PCA were similar to those in Fig. S3 but they also amplified and untangled more clearly the salient population dynamic features in each period. This observation confirms that the orthogonal splitting of sensory and categorical responses is a robust and intrinsic feature of S2, and not an artifact of the chosen set of projection axes. These results are remarkable evidence for the organizing role of S2, in which a tangle of neuronal responses is tidied up at the population level to minimize interference and maximize codification.

### Orthogonal dynamics in S2 associated with the first stimulus and decision-making

To further dissect the low-dimensional orthogonal decoupling between different coding dynamics in a way that is more sensitive to task parameters, we performed a demixed principal component analysis (dPCA). Unlike traditional PCA, dPCA both maximizes the captured variance and effectively separates the population response variability among the different task parameters (5, 7). Applying this approach, we were able to decompose the population dynamics into new dPCs. We performed four kinds of demixing: P1 and P2 stimulus identity, decision, and class. For this, we marginalized neural activity with respect to each one of these parameters and for the four marginalized data we computed the covariance matrices throughout the whole TPDT (−1 to 7.5 s). It is important to stress that the condition independent [see (7)] component was subtracted away from each marginalization. Therefore, the non-coding (or temporal) variance is removed to calculate the demixed axes.

In Figure 4A, we show the population trajectories obtained from the projection over the first two dPCs associated with P1 (Fig. 4A, P1-dPC1 and P1-dPC2). Lines above traces highlight time intervals where P1 can reliably be decoded from single trials (Methods). The population demixed responses merged classes with identical first stimulus, i.e., c1 with c2 (P1 = G, orange and red traces) and c3 with c4 (P1 = E, blue and green traces). Arbitrarily, we chose c2 (orange) and c4 (blue) to be on top in Fig. 4A. In agreement with PCA analysis, P1-dPC1 and P1-dPC2 captured strong sensory and categorical responses (class coding), respectively. Notice that due to the subtraction of the condition independent signal, the P1-dPC1 sensory response displayed a fast fluctuation around zero. Interestingly, the categorical signal (P1-dPC2) encoding P1 identity, re-emerged during P2, arguably for comparison purposes. Thus, the same axis can be used to decode categorical information during the first stimulus and to get information about P1 during the second stimulus. P1-demixed PCA demonstrates how S2 can split encoding dynamics, i.e., by keeping the faithful representation of P1 and its abstract representation into orthogonal subspaces. Furthermore, this different method was able to demonstrate that P1 abstract coding during the first stimulus and its recalling during P2 are both decoupled with the pure sensory dynamics. When we analyzed the neural data marginalized with respect to P2 and identified the most significant axes (P2-dPC 1 & 2), we observed a scenario akin to the one described for P1-dPCs. In this scenario, the dynamics merge classes with identical second stimuli, i.e., c1 with c3 (P2 = G, red and green traces) and c2 with c4 (P2=E, orange and blue traces). The choice of plotting c3 (green) and c4 (blue) on top in Fig. S5 A was totally arbitrary. P2-dPC1 predominantly captured sensory dynamics, which were especially pronounced during the P2 period. Meanwhile, P2-dPC2 revealed a more transformed response during the same task epoch (Fig. S5 A). It’s worth noting that while these findings echo those seen with P1-dPC representations, they provide a sharper understanding of the task.

**Figure 4.**
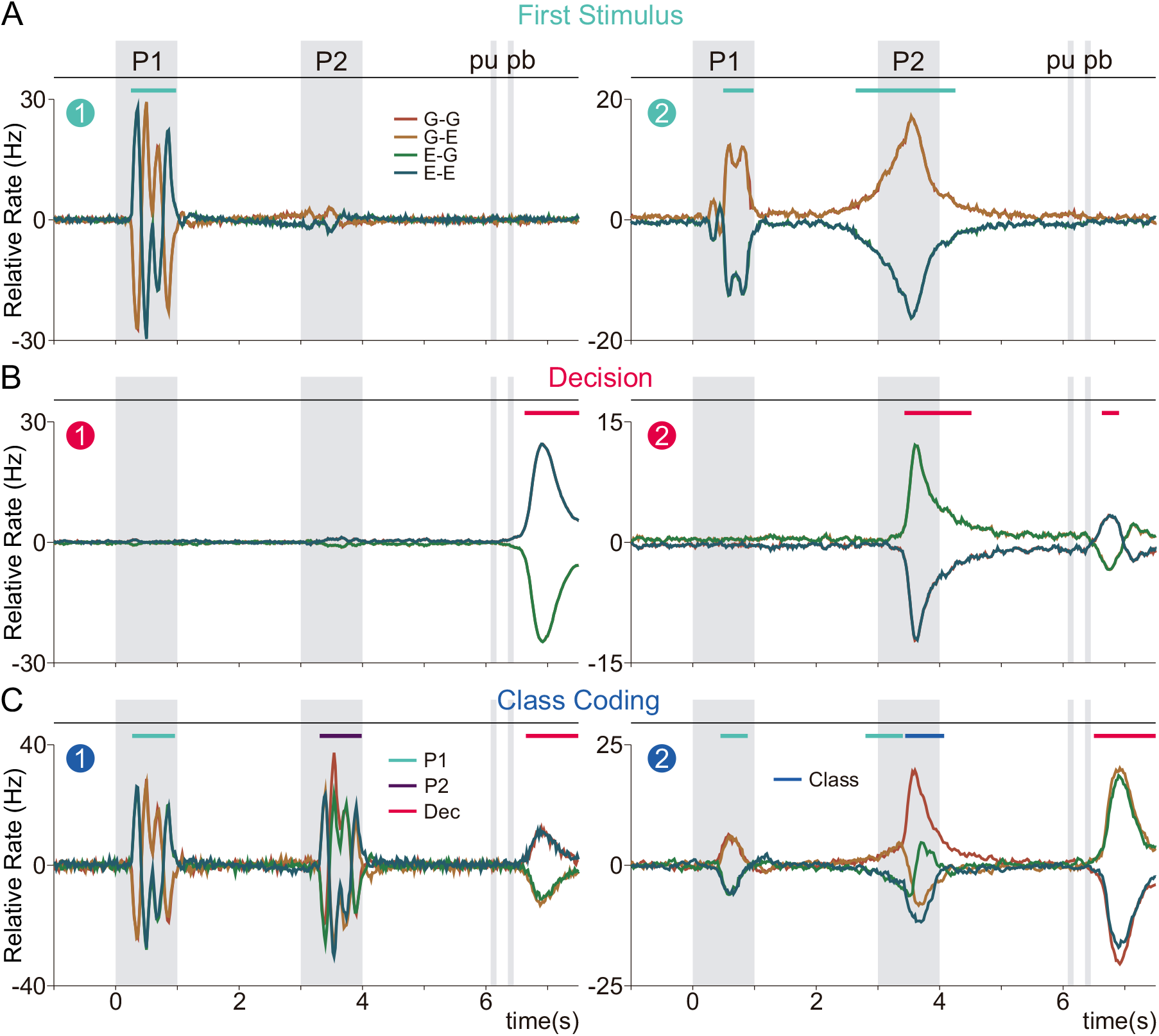
Orthogonal population responses associated with task parameters. Demixed principal component analysis (dPCA) was applied to the marginalized covariance matrices from the whole TPDT (−1 to 7.5 s), with respect to P1 (A) and decision (B), to derive associated demixed principal components (dPCs). Population activity for TDPT (n=1646), sorted by class identity, was projected onto the resulting dPCs axes, and ordered by their EV (P1-dPC1: 6.2%, P1-dPC2: 4.8%; dec-dPC1: 9.1%, dec-dPC2: 4.4%). Only the first two dPCs for both parameters were above the noise level. Color lines on top of each projection represent significant coding periods. With respect to P1, the projections exhibited purely sensory dynamics in the P1-dPC1 and categorical ones in the orthogonal component (P1-dPC2). When projecting onto dec-dPCs, signals purely related to decision outcome emerged after pb (dec-dPC1) and during P2 (dec-dPC2).

Marginalization with respect to decision (Fig.4B), on the other hand, obviously lead to dPCs (dec-dPC1 and dec-dPC2) that merged classes with the same decision outcome (classes matched/non-matched), i.e., c1(G-G) with c4(E-E) and c2 (G-E) with c3 (E-G). Thus, classes c1 and c4 (red and blue traces) as well as c2 and c3 (orange and green traces), overlapped with blue and green lines being on top without premeditation. Projections along decision-dPC1 strictly captured the match (c1 & c4) and non-match (c2 & c3) outcomes after the pb press. This result agrees with the decision coding dynamics observed in Fig. 2A during pb (pink line). We can hypothesize that this inter-trial (after pb) signal might be relevant to learning processes and history dependent changes in the population. Interestingly, dec-dPC2 revealed a decision coding that emerged during the comparison period and remained significant during the first period of the second delay. Notice that the decision variance in Fig. 2A became significant during an analogous period of the task. Notably, the fact that these dynamics exists in orthogonal subspaces would mean, again, that the S2 network employed decoupled population responses to code the decision during P2 and across the inter-trial period. It is important to highlight that to obtain these orthogonal signals it was imperative to apply dPCA to emphasize on the decision dynamics. The applying of PCA to the whole task (Fig. S3) or to a narrow period (Fig. 3), would not have allowed us to visualize these decision responses since they are diluted with the non-coding or temporal dynamics. These results further suggest that S2 employs orthogonality to split decision signals associated with different cognitive processes.

To further elaborate on the previous concept and examine the transition of population response from P1 to decision coding, we investigated the dynamic interaction from each class throughout the task (5). By marginalizing the neural data according to classes (GG, GE, EG, EE), we identified the most significant components (dPCs) related to differences in classes—class-dPC 1 & 2 (Fig. 4C). Consistent with P1-dPCA and dec-dPCA analysis, class-dPC1 predominantly captured sensory dynamics, which were especially pronounced during the P1 and P2 periods. Different classes converge based on the identity of either the first or second stimulus, with traces of varying colors overlapping according to the chosen arbitrary plotting order. On the other hand, class-dPC2 illustrated the divergence of categorical signals, enabling us to observe the interaction of these dynamics within a distinct orthogonal subspace to assist the S2 network in task resolution. Through this analysis, we aim to underline the intrinsic segregation between sensory and categorical signals within the S2, showcasing this dynamic interplay in its clearest form.

In line with Fig. S3 and Fig. 2, the application of dPCA to the LCT population revealed only one significant dPC associated with the stimuli P1 and P2, and class (Fig. S5 B). The resulting dynamics demonstrated sensory responses akin to those depicted for 1st P1-dPC, 1st class-dPC (Fig. 4A & C) and 1st P2-dPC (Fig. S5 A). All traces matched and colors overlapped as previously described. This indicates the absence of categorical signals during LCT. Given that decision coding was not present during LCT, no decision dPCs were significant in this experimental context (Fig. S5 B).

Until now we have provided enough evidence about how the S2 network intertwines the mix of heterogeneous single responses into a complex and organized space where sensory and categorical signals are decoupled and maintained in orthogonal subspaces. Nevertheless, at this point the reader might ask (as we did) if some relation exists between those orthogonal components. In other words, do orthogonal encoding dynamics emerge from a common neuronal substrate or from separate groups of neurons? To initially address this matter, we plotted the neuronal weights from the most significant axes derived by applying PCA (PC1 & 2) or dPCA (P1-dPC1 and class-dPC1 & 2) to the entire TPDT and LCT datasets (Fig. S6 A & B, respectively). We focused on class-dPCs1 & 2, given their ability to vividly capture the division of sensory and categorical dynamics into distinct orthogonal subspaces. We also selected the P1-dPC1 axis due to its relevance in showcasing sensory responses. In TDPT, the weights for both 1st and 2nd PCA and dPCA (class-dPC1 vs class-dPC2 or class-dPC1 vs P1-dPC1) revealed a lack of clusterization in neuronal activity, instead displaying a continuous, roughly uniform distribution. This continuity persisted even when combining axes from different dimensionality reduction methods, such as PC1 vs class-dPC1, which capture the most pronounced sensory dynamics. Importantly, the differences between sensory class-dPC1 and PC1 are consequences of the marginalization with respect to the condition-independent signals (5, 7, 23), which explain most of the network variance (see Fig. 2) and therefore are strongly represented in the PCs but not in the dPCs. Similarly, for the LCT, a continuum of responses was observed when plotting the weights of the most significant axes against one another, reinforcing the absence of any evident clustering in the neuronal responses across both tasks according to linear projection methods.

Up to this point, the results suggest that sensory and categorical dynamics in the S2 network might emerge from a continuous substrate of neuronal activity rather than separate clusters in the TPDT and LCT. Nevertheless, linear methods such PCA and dPCA, may lack the requisite power to discern distinct clusters within the neuronal activity. In light of this, we shifted our analytical focus towards a non-linear method known for its robust capability in identifying clusters (27).

### A continuum of coding responses in S2 underlying orthogonal coding dynamics

To further address this problem, we used a nonlinear dimensionality reduction technique known as Uniform Manifold Approximation and Projection (UMAP) (28). Whereas linear dimensionality reduction methods like PCA and dPCA can only identify the most relevant linear subspaces underlying the organization of a set of *n*-dimension points, UMAP uses differential-geometric ideas to identify linear as well as nonlinear structures, that is topological manifolds, in the projected data. By inspection of the projected data, UMAP can thus be used to study the topology of a set of points and determine the number of connected components, that is, groups or clusters, in which the set is organized. This method was recently used in a variety of contexts, from sorting neuron types (29, 30) to uncovering the toroidal geometry underlying grid cell response (31). Moreover, we applied this non-linear approach to the analysis of DPC population dynamics during a categorization task (1) and for establishing hierarchical responses across the somatosensory sub-areas (32). In this context, it is crucial to emphasize that our application of UMAP in this study is specifically aimed at examining the continuity within the neural substrate and identifying potential clusters of neurons responsible for the observed sensory and categorical dynamics. Unlike linear methods such as PCA and dPCA, UMAP is not and cannot be used to define explicit projection axes. Instead, UMAP’
ss utility lies in its provable ability to detect clusters (or lack of), a capability where linear methods necessarily fall short in general (27).

When we applied UMAP to our S2 population, we either focused on the entire TPDT (−1 to 7.5 seconds, Fig. S7 A) or on ruling out the movement and inter-trial periods (−1 to 6 s, Fig. 5A). Given the high variance observed at the end of each trial (please see Fig. 2), the latter option allowed us to better focus on responses associated with P1, P2 and class coding. To employ UMAP, we mapped each neuron to a point in dimension 4 * *nt*, where *nt* is the number of time points, by concatenating its normalized firing rate vectors in the four classes. We then used UMAP to project the resulting set of points in two dimensions (Fig. 5 & S7). The resulting projections revealed a connected, single-component continuum of responses, with no discrete groups or clusters. To characterize how responses varied across this continuum, we calculated a double-gaussian (σ=0.4) weighted-average of the activity of all neighboring neurons at certain points (colored X-marks) throughout the 2D projection (Fig. 5B & S7B). Although sensory, categorical, and decision response phenotypes can clearly be detected, they all belong to the same continuous deformation of responses across the projection plane. Notice that in the continuum, categorical signals (averages with purple and yellow crosses in Fig. 5A), are surrounded by different kinds of sensory and non-coding responses. Despite these signals are averages in specific locations of the UMAP plane, they are strikingly akin to some of the exemplary neurons shown in Figs. 1, S1 and S2 (some neurons are *b, e, f, h* in Fig. 1, *d, f, n, p* in Fig. S1 and *e, g, l, p* in Fig. S2).

**Figure 5.**
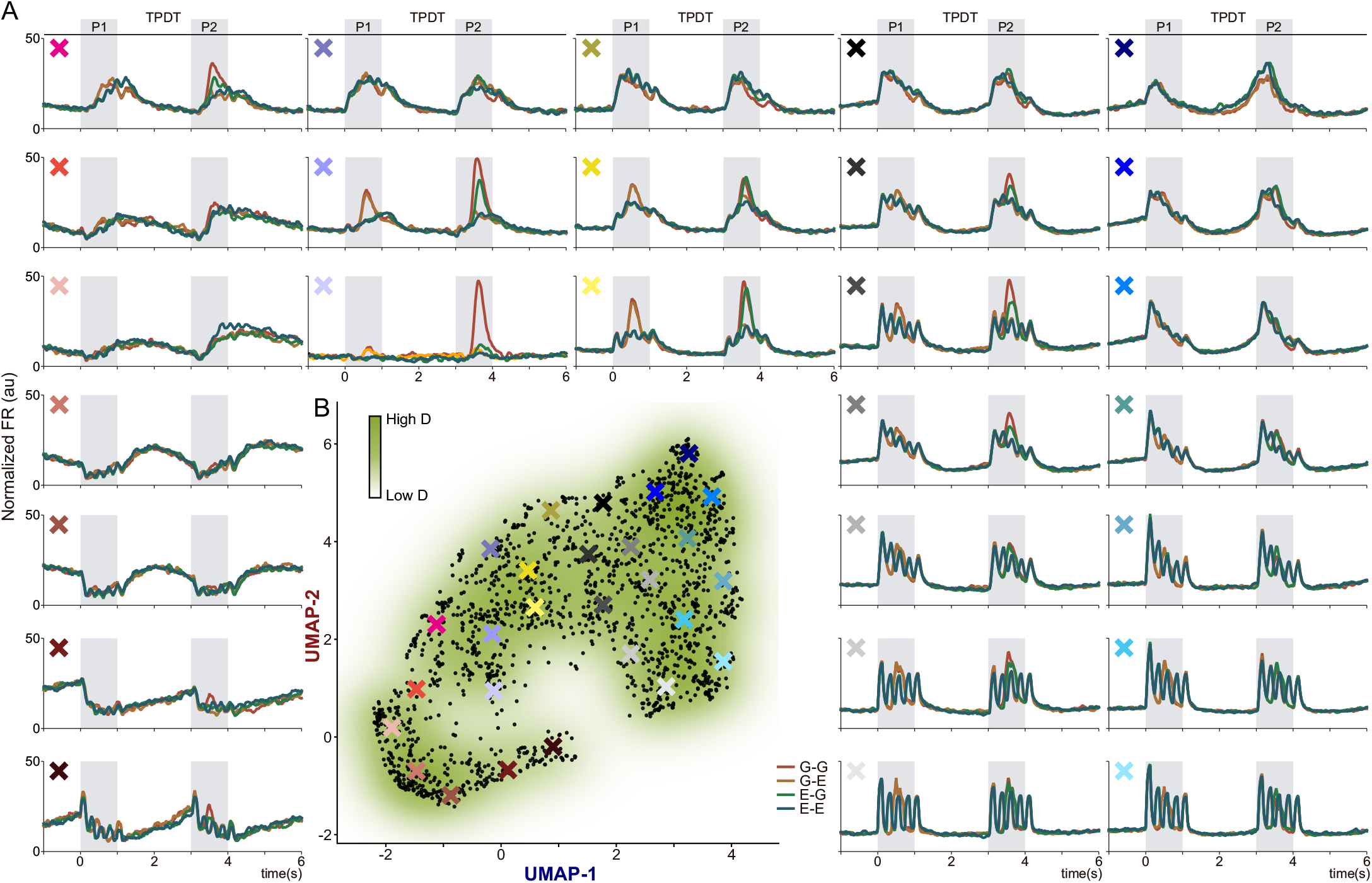
A continuum of responses in S2: from sensory to categorical neurons during the TPDT. To focus on the sensory and categorical responses, we excluded the movement and inter-trial periods. (A) Each subpanel represents a weighted average of neuronal activity and is presented in back-transform arbitrary units that roughly correspond to firing rate values (Hz). Notice that the exemplary averages range from sensory to more categorical and temporal dynamics (bottom, middle and top panels, respectively). (B) 2-D nonlinear UMAP density projection of the normalized activity of all units (n=1646, each black dot) recorded during TPDT (restricted from -1 to 6 s of the task) exhibited a continuum of responses. The color X marks represent the centers of the double-Gaussian (σ = 0.5) weighted averages that are displayed in the panels in A.

Conversely, UMAP projection of neuronal responses in the LCT (Fig. 6 & S8) revealed a disconnected two-component structure where all that remained were decisional and sensory responses, akin to those seen in representative neurons in Fig. S2 (LCT case; some neurons are *a, c, k, m*). The disappearance of the categorical response region is seemingly the main cause underlying the disconnection of the neuronal response manifold in LCT. The contribution of neurons with categorical coding in the S2 network during TPDT thus appears to be crucial for the formation and maintenance of the observed continuum of neuronal responses. Here, we wish to emphasize the distinct advantage of UMAP in identifying context-dependent variations in the continuity of the neural substrate—a task at which linear methods do not (and in general, cannot) perform as effectively, as demonstrated in the scatter plots of neuronal weights in Fig. S6 B.

**Figure 6.**
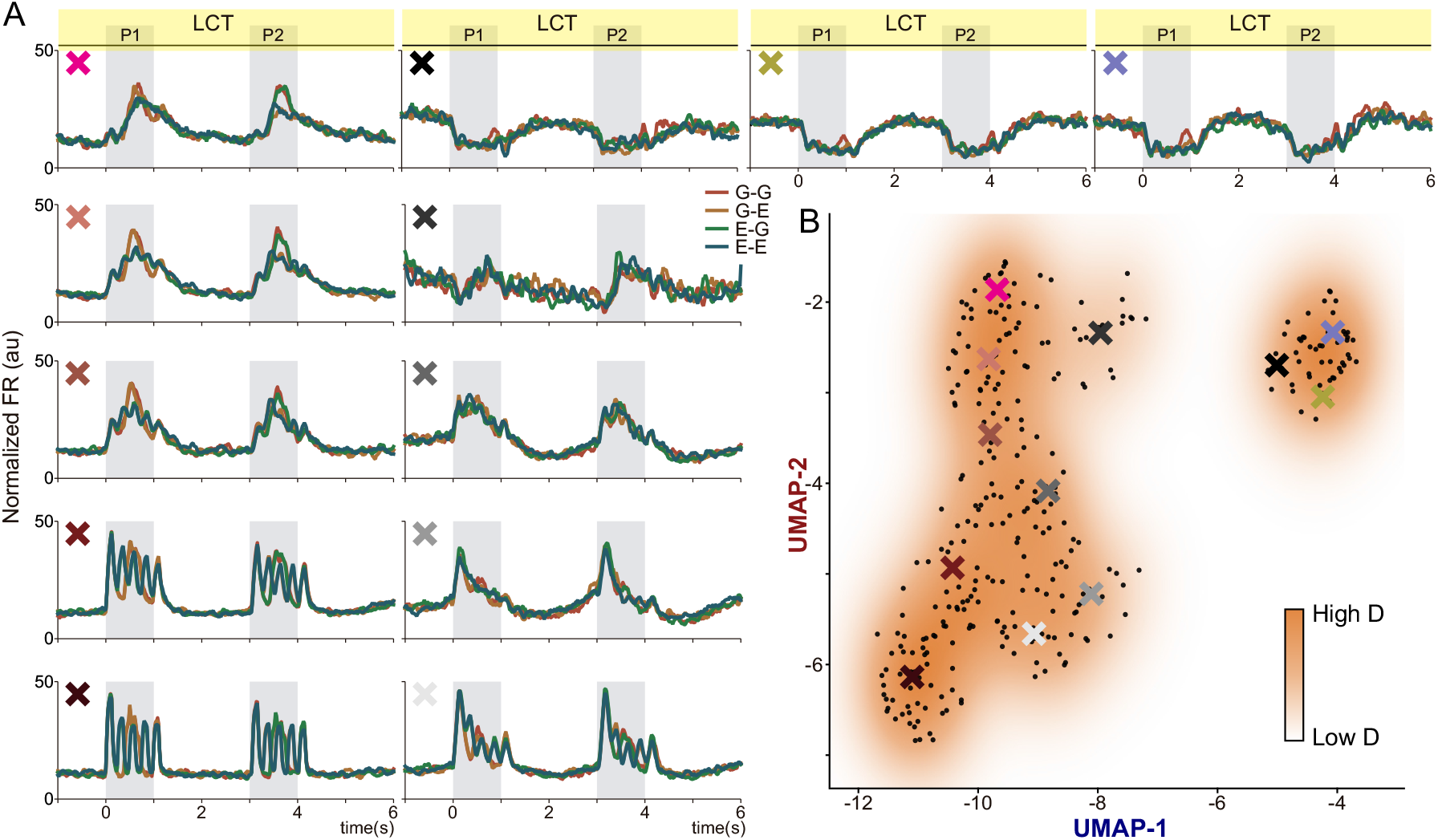
Breaking the continuum in S2 during the non-demanding task. To focus on the sensory and categorical responses, we excluded the movement and inter-trial periods. (A) Each subpanel represents the weighted average of neuronal activity in arbitrary units that approximately correspond to firing rate values (Hz). Given that LCT is a non-demanding task, the neurons that exhibited categorical dynamics during TPDT disappeared, leaving only sensory and temporal representations. (B) UMAP density projection of the normalized activity of all units (n=313, each black dot) recorded during LCT (restricted from -1 to 6 s of the task) showed a breakup in the continuum that emerged during TPDT, exhibiting only sensory and temporal dynamics. The color X marks represents the centers of double-Gaussian (σ = 0.5) weighted averages that are displayed in the panels in A.

In conclusion, UMAP revealed that the orthogonal representation identified by PCA and dPCA analysis from in the S2 population responses emerged from a unique continuous substrate which connects non-coding (temporal), categorical and sensory single unit responses. Notably, the disappearance of categorical signals broke up this substrate into distinct components during LCT. Our work underscores the importance of nonlinear analysis methods to understand the neural encoding of physical stimuli. Going deeper, we asked if there is a network mechanism that could explain the modulation of the neuronal continuum between the TPDT and LCT. We approached this question using qualitative nonlinear dynamical systems modeling of neuronal population activity.

### Feedback gain modulation in a network model of the S2 as a biologically plausible mechanism for the emergence of context-dependent information representation

We used modeling to uncover which mechanisms could explain context-dependent information representation in the S2. We relied on the established fact that the non-linear recurrent neural network (RNN) response is largely determined and can be modulated by the configuration of its eigenmodes and their dynamical properties (33, 34). Eigenmodes are independent directions in the neural state space of a recurrently interconnected neuronal population that can be used to decompose the network dynamics into a sum of characteristic modal responses. Locally around a dynamical equilibrium of the network, eigenmodes are determined by the eigenvectors of linearized dynamics. Each eigenmode is associated with a time constant *τ* = 1/|*λ*| that determines its characteristic timescale of information integration (Fig. S9), where *λ* is the eigenvalue associated with the eigenvector generating the eigenmode. Fast (or faithful) eigenmodes (small *τ*, large negative *λ*) respond rapidly to input and have a short memory, that is, their response converges back to a strongly attractive equilibrium once inputs are removed. Slow (or temporal) eigenmodes (large *τ*, small negative *λ*) integrate inputs slowly and have a long memory, that is, their response largely outlasts input presentation. Finally, metastable/multistable (or categorical) eigenmodes (large *τ*, small positive *λ*) are slow eigenmodes characterized by the existence of a threshold that distinguishes different input categories. The threshold of categorical eigenmodes is determined by the local dynamical instability brought by the small positive mode eigenvalue.

We hand-built (no automatic training used) a nonlinear RNN model with fully tunable eigenmodes (see Methods). The model equations are provided in (Eq. S11). Each node *i* in the network is associated with a state variable *x*_*i*_ representing the overall activity of a neuron (or a neuronal population). A first network interaction matrix *A* models how neurotransmitter liberation affects the node dynamics. In particular, *A*_*ij*_ models the effect of the liberation of neurotransmitter *j* on the activity of node *i*. A second network interaction matrix *B* models how node activation affects neurotransmitter liberation. In particular, *B*_*jk*_ models the effect of the activity of node *k* on the liberation of neurotransmitter *j*. Because of the qualitative nature of our modeling approach, we did not explicitly distinguish between excitatory and inhibitory neurotransmission, a subtlety that will be explored in future works. Finally, gains *k*_*j*_, *j* = 1, …, *N*, model presynaptic modulation: the larger *k*_*j*_, the larger the release of neurotransmitter *j* for the same presynaptic neural activation. With a minor modification to the model, the gains *k*_*j*_, *j* = 1,…, *N*, can equivalently model postsynaptic modulation, i.e., the larger *k*_*j*_, the larger the effect that neurotransmitter release has on the activity of neuron (or neuronal population) *j*.

By tuning the strength of recurrent interconnections, gains *k*_*j*_, *j* = 1,…, *N*, are feedback gains that shape the dynamics and thus the information-processing of the network eigenmodes. In particular, by controlling the strength of recurrent amplification, these feedback gains control the local stability of the network eigenmodes, which in turn determines the mode encoding properties: a small feedback gain is associated with highly stable (fast/faithful) eigenmodes; a larger feedback gain is associated with weakly stable (temporal) or metastable/multistable (categorical) eigenmodes.

We hand-built a family of networks with a mixture of faithful, temporal, and categorical eigenmodes and tested their response to extended or grouped stimuli. The obtained responses were then merged into a metapopulation for analysis, as done in experiments. To introduce heterogeneity and variability, we generated faithful, temporal, and categorical eigenmodes by sampling the associated feedback gains according to distributions centered around a different characteristic value *k*_0_ for each mode type (Fig. 7A). The components of the eigenvector associated with each eigenmode determined the selectivity (pure or mixed) of each modeled neuron. The final network consisted of a mixture of neurons with either pure (one third) or mixed (two third) selectivity (see Methods for details). Of the neurons with pure selectivity, half were purely temporal and half purely categorical. Neurons with mixed selectivity varied between predominantly faithful and predominantly categorical. Mixed selectivity between temporal and faithful and between temporal and sensory responses was not enforced because part of the faithful neurons, those associated with larger feedback gain, and part of the categorical neurons, those associated with smaller feedback gain, already exhibited purely temporal responses.

**Figure 7.**
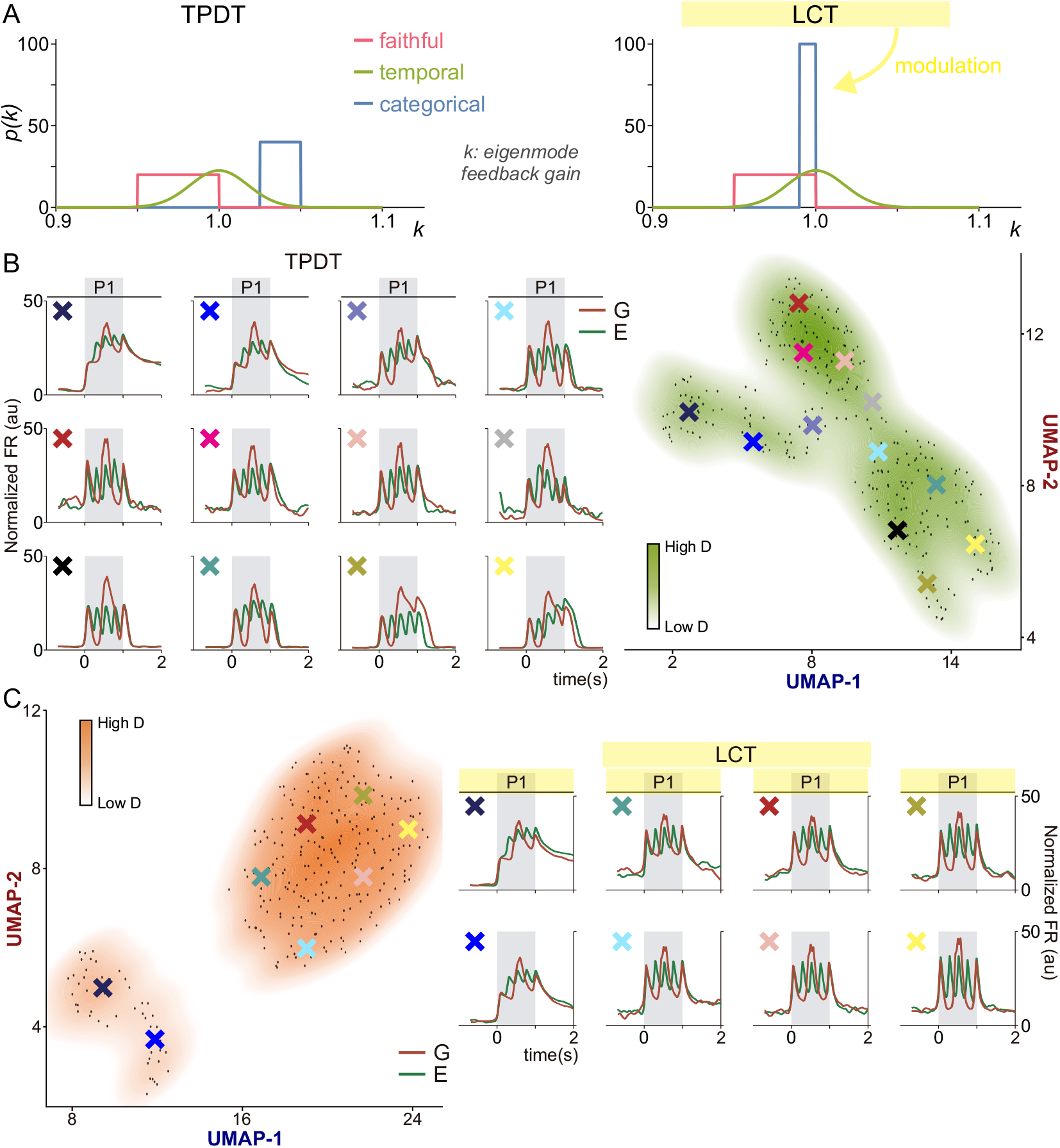
A network model of the S2 neuronal population dynamics. The network reproduces the transition from a continuum of responses in TDPT to clustered responses in LCT. (A) Faithful, temporal, and categorical eigenmodes in our network were defined by different characteristic network feedback gains (see main text and Methods). For each eigenmode type, the defining feedback gain was sampled from the distribution sketched in the panel to create a family of heterogeneous eigenmodes over which the network structure was built resulting in a mixture of pure and mixed neurons. The transition from active to control was modeled by modulating the feedback gain distribution of the categorical eigenmodes as drawn. (B) Similar to Figure 5 but for the network model. In active condition, the modeled S2 neural manifold exhibited a continuum of responses connecting faithful, temporal, and categorical modes. (C) Similar to Figure 6 but for the network model. In LCT condition, categorical responses disappeared from the modeled S2 neural manifold, which led to its break-up into two sharply separated clusters of purely temporal and faithful responses.

To model the transition from active (TPDT) to control (LCT) context as observed in experimental data (Fig. S2), we shifted the distribution of feedback gains associated with categorical representations toward smaller values and overlapped it both with sensory and temporal feedback gain distributions (Fig. 7A), thus letting categorical representation effectively disappear from the modeled population dynamics. When we ran UMAP over the generated neuronal population in the active and control contexts we obtained the projections in Figure 7B. Similarly to what obtained on the S2 neuronal population (Figures 5,6 and Suppl. Figures 7,8), in the active context the simulated neuronal population activity was distributed along a connected manifold forming a continuum of activity types connecting purely faithful, temporal, and categorical responses. Conversely, in control context the population activity manifold broke into two well separated clusters due to the disappearance of the categorical response. It is important to stress that the breaking of the response continuum in control was not enforced ad-hoc. Indeed, the feedback gain distributions determining the eigenmode family were more contiguous in the modeled control context as compared to active. It is the functional disappearance of the categorical response, bridging purely faithful to purely temporal responses in the active context, that breaks the response continuum in control.

Our model suggests a simple and general biological mechanism to explain the emergence of context-dependent information representation in the S2, namely, modulation of the feedback gains determining the strength of recurrent interconnection-mediated nonlinear amplification through pre- or postsynaptic modulation. More specifically, a neuromodulation-mediated decrease of the feedback gain of categorical eigenmodes is sufficient to remove the local dynamical instability determining their categorization threshold and transform them into either temporal or faithful eigenmodes. Both pre- and postsynaptic modulation are plausible biological candidates to realize such a feedback modulation mechanism. Both in biology and in modeling, the disappearance of categorical eigenmodes is the key factor determining the breaking of the response continuum.

## Discussion

Our research provides five critical insights into the functions and capabilities of the S2 neuronal population. First, in line with our previous studies, neurons in S2 displayed a multifaceted diversity of responses, characterized by a palette of faithful (sensory), categorical, and temporal signals. This diversity of neural substrates positions S2 as a key driver in sensory abstraction, possibly serving as a primary source of inputs for downstream hierarchical processes. Such observable heterogeneity and mixed selectivity also suggests S2’s potential for highly efficient computations (35–37). Second, using two different contexts distinguished by their cognitive demand revealed that S2 context sensitivity is different as compared to other areas. Unlike in area 3b of S1, where only faithful and context-invariant responses are recorded (20), or in DPC, where the context dependent coding primarily aims at comparing stimuli and forming decision-related responses, S2 displayed neurons with combined context-dependent and context-independent coding. While this coding versatility may partially stem from the array of responses mentioned earlier, it also hints at underlying complexities in S2 that go beyond single unit response cataloging. When the task requires, S2 possesses the capability for managing and segregating diverse signals, but during a non-demanding task this processing is turned off. Third, our population analysis not only corroborated this hypothesis but also unveiled a mechanism within the S2 network through which sensory and transformed dynamics are separated into orthogonal subspaces. Interestingly, this arrangement of signal splitting was observed in both brain hemispheres, with minor yet not significant differences in their lateralization relative to the stimulation. While similar mechanisms have been identified in other neuronal populations (2, 4, 6), we believe our findings present a network uniquely designed to differentiate between sensory and categorical dynamics. Fourth, the use of a nonlinear dimensionality reduction method tailored to identifying clusters (or lack of) in high-dimensional data structures, revealed that these dynamics arise from a context-dependent continuum of neural activity, rather than from isolated neuronal clusters. Thus, the identified orthogonal representation directions emerged from a single continuous substrate, where sensory, temporal (non-coding), and categorical responses are smoothly interconnected. Fifth, a network model suggested that the context-dependent differences observed in S2 population dynamics can be explained through a simple feedback gain-modulation mechanism. This observation provides well-known attentional mechanisms in the brain (38, 39) with a new mode of action, namely, switching on and off specific coding dynamics and shaping the neural manifold where neuronal population coding happens as a function of context.

The question of whether the brain actually employs orthogonality for dissociating signals at the population level or whether the evidence for orthogonal representations is due to the used dimensionality reduction techniques (40) has been a topic of rigorous discussion (3, 41). A recent study has shown that the orthogonal dynamics associated with sensory and mnemonic coding appeared during implicit task learning (4). Their results in (4) suggest that orthogonal population dynamics are not present in naive animals, but they emerge through familiarization with the task. They also show that this mechanism might have its roots in individual neurons, specifically in those that either maintain (‘stable neurons’) or reverse (‘switching neurons’) their selectivity over time. The authors of that study found that the interplay between these distinct neuronal dynamics enables the rotation of population representations. This allows a unified network of neurons to effectively encode both sensory data and short-term memories, thereby optimizing computational efficiency. In S2, the decoupling of sensory and categorical dynamics into well-defined subspaces might have its origin in neurons which purely encode the different signals but also in those that exhibit mix dynamics which probably could mingle their level of coding between sensory and categorical. Thus, the S2 network would be able to dynamically dissociate what has already been transformed from the raw sensory input, reducing interference, and maximizing flexible computations. To support this finding, we applied period-restricted and whole task approaches in addition to parameter marginalized dimensionality reduction analysis. We found similar results with all these different approaches, suggesting that the orthogonal separation of the dynamics is an intrinsic feature of the S2 network, rather than an artifact arising from the selected projection axes.

Importantly, the emergence of the coding orthogonal subspaces may be related to the underlying network communication structure. Prior research proposed that the development of orthogonal, decoupled dynamics may act as ‘communication subspaces’, serving as an effective framework for enhancing inter-area neural communication (42). In their study, the authors showed that activity fluctuations in the secondary visual area (V2) are associated with a specific subset of activity patterns in the primary visual area (V1). Our work builds upon the strength of this study too, given that the uncoupled subspaces that emerged from our analysis could play a role to enhance communication with other brain areas. For instance, one might guess that sensory signals are involved in communication with areas of S1, categorical signals with frontal and premotor cortices, and push-button ones with the execution of decisions by motor cortices. Further research employing simultaneous recording among several areas is required to clarify the role of these orthogonal dynamics for communicating signals across the brain in the task employed here.

An intriguing observation that emerged from our population analysis is that orthogonal coding might also be used by the S2 network to differentiate decision signals. While there was significant decision-related coding observed both before and after push button movement (inter-trial period), our dPCA results reveal that these two decision signals are also orthogonally arranged. This suggests that they may be involved in distinct functions. For instance, coding during the second stimulus may be related to the comparison, whereas post-movement activity might pertain to evaluating the reward associated with the decision (43). This means that S2 responses could be influenced by decision history, hinting at a more nuanced function for this area than previously suggested. Such a mechanism might form part of a feedback loop that continually refines the network’s performance based on recent experiences. This process may also be facilitated by the unique continuum substrate of responses upon which the network is based. This opens an interesting hypothesis to be verified in future experiments studying the changes in S2 dynamics during learning.

Our last hypothesis opens, in turn, an intriguing question: What are the potential benefits of a network built upon a continuous substrate of responses? Might such a feature promote the development of orthogonal subspaces? To investigate the continuity of the neuronal encoding substrate, we employed UMAP. In the 2D visualization, distinct patterns in averaged neural activity were evident (Fig. 5), with sensory and temporal signals flanking the categorical ones. Yet, density-based clustering on these UMAP projections did not identify any clear subpopulations. The property of the S2 network of having ordered but continuously connected patches of neuronal activity types is in line with recent findings where orthogonal encoding strategies were also uncovered. Recent research, where monkeys were trained to memorize sequential spatial locations of three visual stimuli, discovered that the frontal network encoded each stimulus position into a distinct orthogonal subspace (6). The authors analyzed the anatomical and functional organization of individual neurons and found, similar to us, a family of activity patches intertwined within a unified substrate. They posit that such spatial arrangement might offer advantages in positive feedback learning, coordinate transformations, and even in integrating multiple sensory modalities. Even if our recordings preclude spatial analysis of S2’s neurons, we hypothesize that the lack of functional clusters may also be related to a structural continuity. We propose this line of study for future exploration.

The use of RNNs has emerged as a groundbreaking tool in cognitive neuroscience, offering deep insights into the mechanisms hidden within neural dynamics (12, 34, 44–46). Like many studies referenced here, our aim was to design an artificial model that mirrors the dynamics triggered by the initial stimulus in our task. By adjusting the strength of positive feedback gains, we successfully shaped the coding dynamics based on context. However, as the reader may also ask: how might this mechanism operate within a biological substrate as the S2 network? One hypothesis is drawn from local field potentials (LFPs). In a visual task, it was observed that distinct propagation directions of alpha and gamma oscillations provide an efficient feedback and feedforward control mechanism that regulates information transmission (47). In that research, the authors discovered that stimulation of the deeper layers prompted the spread of alpha oscillations to V1, dampening activity in this region. This gain modulation associated with LFP bands has been also suggested by subsequent studies (48). Parallel research in a somatosensory frequency discrimination task has also explored the behavior of alpha oscillations across cortical areas (49). The study found that lower alpha power correlated with increased firing rates in several areas, including S2. Particularly, S2 exhibited an increase in alpha power during the beginning of the working memory period which corresponded with a decrease in activity and coding in this network. This means that the modulation in alpha power might serve as a feedback mechanism, facilitated by higher-order areas, optimizing the processing of relevant sensory data. While in this study we did not center on examining oscillatory activity in S2, the findings presented thus far open promising avenues for future research within this task.

In sum, our exploration into the intricacies of S2 has unveiled a dynamic landscape of neural responses, adaptive behaviors associated with the S2 population responses, and shows an innovative mechanism underpinning sensory abstraction and processing. The discovery of an intrinsic coding orthogonalization strategy, the intricate continuum of neural activity, and the nuanced interplay of decision-related signals all attest to the rich complexity of S2. As we build upon prior findings and integrate novel methodologies like RNNs, we are not only charting new territories in our understanding of neural dynamics but also shaping the trajectory of future neuroscience research. The intricacies of S2, as showcased, hold profound implications, offering fresh perspectives on the brain’s computational prowess and adaptability.

## ACKNOWLEDGMENTS

We thank Hector Diaz for his technical assistance. This work was supported by grants PAPIIT-IN205022 from the Dirección de Asuntos del Personal Académico de la Universidad Nacional Autónoma de México (to R.R.-P.) and CONAHCYT-319347 (to R.R.-P.) from Consejo Nacional de Ciencia y Tecnología; IBRO Early Career Award 2022 (to R.R.-P.) from International Brain Research Association. L.B. is a postdoctoral student (Postdoctoral fellowship CONACYT-838783).

## Methods

### Temporal pattern discrimination task (TPDT)

The TPDT used here has been previously described (17, 20). Briefly, two monkeys (*Macaca mulatta*) were trained to report whether the temporal structure of two vibrotactile stimuli patterns (P1 and P2) of equal mean frequency (5 Hz, 5 pulses) were the same (P2 = P1) or different (P2 ≠ P1; Fig. 1A). The temporal structure of each pattern was either grouped (G) or extended (E) with a fixed stimulation period of 1 s. The five pulses were delivered periodically during the extended pattern (E), and three grouped centered pulses with a smaller distance between them as compared to the first and final pulses, during the grouped pattern (G). Monkeys performed the task in blocks of trials in which the two stimulus patterns had a fixed mean frequency. The right arm, hand and fingers were held comfortably but firmly throughout the experiments. The left hand operated an immovable key (elbow at ∼90°) and two push buttons in front of the animal, 25 cm away from the shoulder, at eye level. Stimuli were delivered to the skin of one digit from the distal segment of the right restrained hand, via a computer-controlled stimulator (2 mm round tip, BME Systems, Baltimore, MD). The descending of the probe until achieves a skin indentation of 500 μm, marked the beginning of the trial (probe down event, “pd” in Fig. 1A). After pd the animals gently grabbed with their free left hand an immovable key (key down event, “kd” in Fig. 1A). Vibrotactile stimuli consisted of trains of short mechanical pulses; each pulse consisted of a single-cycle sinusoid lasting 20 ms. Time is always referenced to first stimulus onset (0 s corresponds to the start of P1). In a trial, P1 and P2 were delivered consecutively to the glabrous skin of one fingertip, separated by a fixed inter-stimulus delay period of 2 s (1 to 3 s). Each stimulus could be one of the two possible patterns: grouped (G, upper trace of Fig. 1A) or extended (E, lower trace of Fig. 1A) pulses. Therefore, in total there were four possible P1-P2 combinations, denominated as classes: G-G (class 1, c1), G-E (class 2, c2), E-G (class 3, c3) and E-E (class 4, c4). These were presented in pseudo-random order to the monkeys across trials. The monkeys were asked to report whether P2 = P1 (match: combinations E-E and G-G) or P2 ≠ P1 (non-match: combinations E-G and G-E) after a fixed delay period of 2 s (4 to 6 s) between the end of P2 and the mechanical probe rising from the skin (probe up event, “pu” in Fig. 1A). The “pu” functioned as a go signal that triggered the animal’s release of the key (key up event, “ku” in Fig. 1a). The monkey indicated their decision by pressing one of two push buttons with the left hand (push button event, “pb” in Fig. 1A, lateral push button for P2 = P1, medial push button for P2 ≠ P1). As the two stimulus patterns had equal mean frequency over their full duration (1 s), the decision had to be based on comparison of their temporal structure. The animal was rewarded for correct decisions with a drop of liquid. Animals were handled in accordance with standards of the National Institutes of Health (NIH) and Society for Neuroscience (SfN). All protocols were approved by the Institutional Animal Care and Use Committee of the Instituto de Fisiología Celular (IFC), Universidad Nacional Autónoma de México (UNAM).

### Light control task (LCT)

In this task variant, events proceeded exactly as described above and in Fig. 1A, except that when the probe touched the skin (“pd”), one of the two push buttons was illuminated, indicating the correct choice throughout the trial until pb. Identical stimuli were used. The monkey grasped the key until the probe was lifted, but in this case the light was turned off when the probe lifted from the skin. The monkey was rewarded for pressing the illuminated button. Maintaining stimuli and arm movements identical to the TPDT, the decision must be based on the visual stimuli instead.

### Task design and performance

The TPDT is not a simple variation of the vibrotactile frequency discrimination task (VFDT) (50). Some cognitive demands and the basic structure of the tasks are similar: both require attention to two separate vibrotactile stimuli (TPDT: P1, P2; VFDT: f1, f2), working memory and a comparison to reach the decision report. Nevertheless, the TPDT requires a very different evaluation of the stimuli; as they only differ by their temporal structure, any computation must be restricted to the internal structure to identify, categorize, and distinguish between them (20). Further, the comparison process is significantly different between the two tasks. Expanding on the necessitated computation, the VFDT can be solved by computing a difference between the parametric representation of the stimulus frequencies to indicate whether f1 > f2 or f1 < f2, whereas the TPDT offers no comparable method of solution (in any trial P1 and P2 always have the same mean frequency). The TPDT demands a match (P2 = P1) vs. non-match decision (P2 ≠ P1). Hence, the comparison employs categorical representations (instead of parametric) of the stimulus patterns. We computed the average performance across S2 recording sessions (p = 84.0%; Monkey RR17, p = 84.5% and Monkey RR20, p = 83.1%). Average performance for TPDT, LCT and each class, is shown in Fig. 1B. Although each animal received around two years of training, this task was difficult enough to impede 100% performance; this reflects the very high-cognitive demands of the TPDT. To provide some context, the average training period to achieve similar performance levels for the VFDT was about six to eight months; for the vibrotactile detection task, the average time was two months (50). After training in the TPDT, the monkeys saturated their average performance around 84% (Fig. 1B, n_SES_ = 423 recording sessions; Monkey RR17, n_SES_ = 281; Monkey RR20, n_SES_ = 142). In addition, the performance was statistically identical for each class (20). Notably, task repetition across recording sessions did not improve performance. However, the performance for the LCT was consistently 100% (Fig. 1B, n_SES_ = 76 recording sessions; Monkey RR17, n_SES_ = 49; Monkey RR20, n_SES_ = 27); this reflects the lack of cognitive demand required for the guided task, as intended by design. As a final observation, the animals were first trained in the LCT, and then gradually introduced to the TPDT. During the recording sessions in S2 (Fig. 1C), animals switched between performing the TPDT and the LCT.

### Recordings

Neuronal recordings were obtained with an array of seven independent, movable microelectrodes (2–3 MΩ) inserted into S2 (Fig.1 C), either contralateral (left hemisphere) or ipsilateral (right hemisphere) to the stimulated hand. We were careful to record just above the primary auditory cortex (A1), and we tested this by using auditory stimulation to ensure that neurons were only responding to vibrotactile stimuli (17–19). The receptive fields of the neurons recorded were all very large and some were bimanual, and since the monkey’
ss hand was carefully fixed in the same manner during each recording session, we do not believe it is possible for these neurons to be responding to motor data in a categorical manner, as would be seen in the parietal ventral area (PV). Coherently, categorical decision responses during P2 or after pb, disappeared during the LCT (see Fig. S2). We recorded 20 trials per stimulus pair (c1; c2; c3; c4). Recording sites changed within terms of recording sessions where one term lasted approximately 30 sessions (1 day = 1 session); the locations of the penetrations were used to construct surface maps in S2 by marking the edges of the small chamber (7 mm in diameter). It is important to emphasize that neurons which displayed sensory or transformed responses were recorded across the entire S2 region, and the possibility of identifying subgroups responses is extremely low (see Fig. 5 for continuum of responses). The neuronal recording protocol was identical for both the TPDT and LCT.

### Datasets

We recorded 1646 S2 neurons using the TPDT, with the stimulus set with 5 Hz mean frequency (Monkey RR17, n=1035; Monkey RR20, n=611). In addition, 313 neurons (Monkey RR17, n=189; Monkey RR20, n=124) were recorded sequentially with TPDT and LCT, with also the 5 Hz mean frequency set. For each neuron of the datasets (n=1646 and n=313), we calculated a time-dependent firing rate per trial using a 100 ms deterministic square kernel with 50 ms steps, beginning 1 s before stimulus pattern P1 and continuing until the end of the trial (1.5 s after the push button press). In previous work we have seen that this window-width is optimal for decoding pattern information in this same area (Fig. S10 in (17)). Importantly, each dataset is defined by four dimensions: N, number of neurons; C, stimulus conditions (classes, always 4); T, time (−1 to 7.5 s, always 170 bins); K, number of hit trials (for each class). A remarkable feature of this task design is the low number of stimulus conditions (four classes), which were equally demanding for the subject. This design allowed us to have, on average, 15.3 hit trials (and 2.9 error trials) per stimulus class for each studied neuron.

### Population Analysis

Single S2 neurons displayed a large repertoire of neuronal responses associated with one or several components of the TPDT (see Fig. 1, S1 & S2). Here, we focus our analysis on the temporal neuronal population signals. For each neuron, we averaged per class the time-dependent firing rate of hit trials (c1, c2, c3 or c4). Using the peri-stimulus time histogram (PSTH) of each neuron, we constructed pseudo-simultaneous population responses by combining neural data mostly recorded separately. For each time and class, the population response is defined by an N-dimensional vector in which each component represents the firing rate from a different neuron. This means that including all the recorded neurons (n=1646), we obtained a 1646-dimensional firing rate vector that depended on the time and class 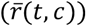. The population firing rates averaged over all hit trials 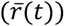 was an N-dimensional vector that measures the mean response for each neuron (*r*^*i*^(*t*)) as a function of time. For the LCT, the control condition, the population response was a 313-dimensional firing rate vector.

In the next equations, we employed the following notation: 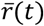 is the firing rate average over all hit trials at each time bin (joining the four classes, c1-c4), 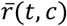 is the firing rate average per class (grouping trials according to each of the four classes), 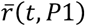 is the firing rate average per P1 stimuli (splitting trials according to P1-identity), 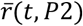 is the firing rate average per P2 stimuli (separating trials according to P2-identity) and 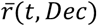 is the firing rate average per decision (dividing trials according to decision). To further explain this measure, the *i* component, *r*^*i*^(*t*), represents the firing rate average of neuron *i* at time *t*, across all trials. Similarly, *r*^*i*^(*t, Dec*) represent the firing rate average of neuron *i* at time *t* across trials with the same decision outcome (P2=P1 or P2≠P1).

### Instantaneous Coding Variances across the population

At each time point, the population instantaneous coding variance (*Var*_*COD*_, Figs. 2A and 2B, blue trace) was computed as the quadratic square sum of the firing rate fluctuations among classes and neurons:

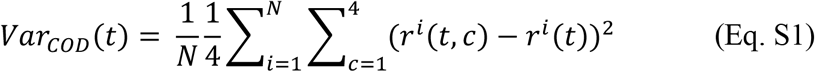

This metric, normalized per neuron, measures the population’s variation of firing rate between classes at each time bin. With this definition, variation will be due to any class-related change in the population activity and due to any residual noise.

To evaluate the influence of each kind of coding, we extended this metric to calculate the instantaneous variance associated with each specific task parameter. At each time bin, the population instantaneous P1 variance (*Var*_*P1*_, Figs. 2A and 2B, light blue trace) was computed as the quadratic square sum of the firing rate fluctuations among P1 identity and neurons:

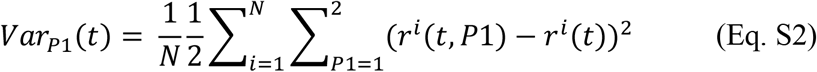

Analogously, the population instantaneous P2 variance (*Var*_*P2*_, Figs. 2A and 2B, purple trace) measures the firing rate fluctuations among P2 identity and neurons:

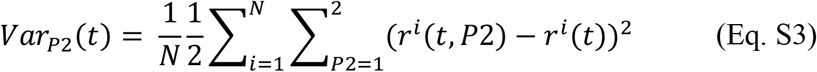

The population instantaneous decision variance (*Var*_*Dec*_, Figs. 2A and 2B, pink trace) measures the firing rate fluctuations among decision identity and neurons:

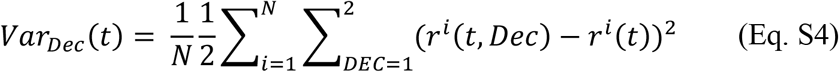

The value of *Var*_*COD*_ during the period immediately before P1 onset represented the inherent residual noise in the firing rate estimates (∼2 sp/s); to be interpreted as a degree of population coding, *Var*_*COD*_ should be higher than this resting state variance (basal variance). The same reasoning applies to the other specific variances. To facilitate the comparison across parameters, values associated to basal variance were subtracted in each kind of variance computed.

### Instantaneous Temporal Variances across the Population

At each time point, the population instantaneous temporal variance (*Var*_*Temp*_, Fig. 2C and Fig. 2D) with respect to the mean firing rate, was computed as the quadratic square sum between the mean firing rate for each time bin (*r*^*i*^(*t*)) and the mean firing rate across the whole trial *r*^*i*^ (from -1s to 7.5s) among neurons:

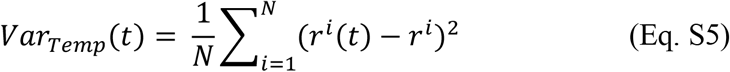

### Principal Component Analysis (PCA)

The main aim of PCA is to find a new coordinate system in which the data can be represented in a more succinct and compact manner. In other words, the idea is to define a low-dimensional subspace that captures most of the variance of the high-dimensional neural space. To characterize how the population activity covaries across classes as a function of time, we performed PCA (Figs. 3 & S3) over classes (c1, c2, c3 or c4) where we combined variance over classes and time (from -1s to 7.5s in Fig. S3 and smaller periods in Fig. 3). PCA yields a new coordinate system for the N-dimensional data, in which the first coordinate accounts for as much of the variance of the neural population. The second coordinate accounts for as much of the remaining variance, and so on; however, each subsequent axis is restricted to be orthogonal to all previous axes. PCA was computed from the firing rate covariance matrix, that averages over time bins, *t*, and classes, *c*:

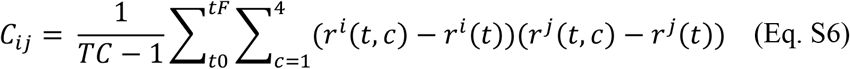

where *T* denotes the number of time bins between *tF* and *t0*, and *C* is the number of classes (4 in our task), *r*^*i*^(*t, c*) denotes the trial-averaged firing rate of the neuron *i*, under class *c*, at time *t*, and *r*^*i*^(*t*) is the firing rate average of neuron *i* across classes at time *t*. In Fig. 3 and S3, PCs were calculated for TPDT (n=1646) and LCT (n=313). The diagonalization of the covariance matrix, *C* = *UDU*^*T*^, yields a new coordinate system given by the columns of the matrix *U*. We refer to the columns of *U* as the principal components (PCs). On the other hand, *D* is a diagonal matrix of positive values. The diagonal elements of *D* give the amounts of population activity variance captured by the corresponding PCs. We then ordered the PCs depending on this amount of variance captured. The projection of the N-dimensional data onto the *k*th PC is given by:

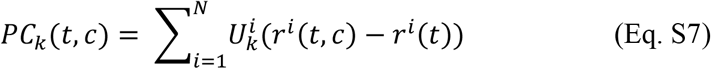

where 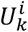 is the *i* element of the *k* PC (*U*_*k*_). Therefore, the PCs are linear readouts of the population activity; in other words, they are linear combinations of the firing rates of the individual neurons. Thus, the contribution of each neuron to a given *PC*_*k*_ is given by the *i*th element of *U*_*k*_. These PCs can be thought of as a low-dimensional description of the population activity in this coding subspace.

The time-windows for P1 and P2 (0.6 s) and decision (1.5 s post “pb” event) in Fig. 3 A, B, and C were based on our prior findings (Figure 3 in Rossi-Pool et al., 2021b), which utilized ROC analysis to assess the coding capacity of S2 neurons. This identified the optimal periods where S2 neurons (n = 1646) showed the highest coding for those task parameters during TPDT. These periods coincided with a peak in relevant coding and were thus chosen for the narrowed-PCA analysis.

### Demixed Principal Component Analysis (dPCA)

The algorithmic details and mathematical justification for this method were outlined in (7). The method has a supervised and unsupervised part. In brief, dPCA decomposes the neural activity by different chosen task variables to compute marginalized covariance matrices (this is the supervised part, similar to choosing the variables to fit a linear model). Afterward, it carries out a principal component-like analysis over those matrices (this is completely unsupervised). In our task, we marginalized the population activity (*X*) with respect to: P1 identity 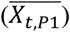 identity 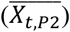, class 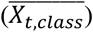 and decision outcome 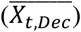. To calculate the marginalization averages,we use the N-dimensional population activity:

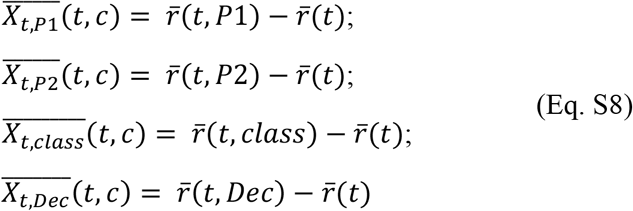

Then, 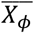 denotes the marginalized matrices with *ϕ* ∈ {{*t, P*1}, {*t, Dec*}} (all the marginalization variables used in this study). Further, 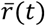 is the firing rate average over all hit trials at each time bin (joining the four classes, c1-c4), 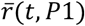 is the firing rate average per P1 stimuli (splitting trials according to P1 identity, G or E), 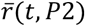 is the firing rate average per P2 stimuli (dividing trials according to P2 identity, G or E), 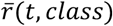 is the firing rate average per class (separating trials according to the four different classes or conditions, c1: G-G; c2:G-E; c3: E-G; c4: E-E), and 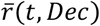 is the firing rate average per decision (dividing trials according to decision: P2 = P1 or P2 -=I P1). All 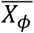 have the same sizes (7). Once marginalization is performed, dPCA finds separate decoder and encoder matrices for each -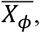 by minimizing with reduced-ranked regression the term:

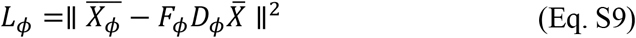

where 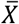 is the centered whole population data matrix (i.e., the average activity of each neuron is 0). The solution of this problem can be obtained analytically in terms of singular value decompositions. Each component *D*_*ϕ*_ can be ordered by the amount of explained variance. The most prominent decoding axis is called the 1^st^ demixed principal component (1^st^ dPC or dPC1) of variable *ϕ*. To avoid dPCA over fitting, we introduced a regularization term and performed cross-validation to choose the regularization parameter. To obtain Fig. 4A & B, we projected the N-dimensional data for a given class 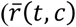, this is the firing rate average per class computed by grouping trials according to each of the four classes) onto the most prominent decoding axis (*k*) of a given variable *ϕ*. These projections were given by:

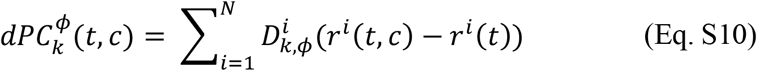

where 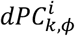 is the *i* component of the *k* most relevant axis of a demixed variable *ϕ*. Here we emphasize that dPCA finds the decoder and encoder of each marginalized variable, *ϕ*, minimizing each *L*_*ϕ*_ separately. This means that we can calculate individually the demixed decoders for each parameter: P1-identity, P2-identity, class, and decision outcome. Figs. 4 A, B & C as well as Fig. S5 A, shows the most prominent demixed components for P1, decision, class and P2, respectively. This is also valid for the analysis in Fig. S5 B where the projections were performed over the most significant dPCs obtained from the marginalized variables P1, P2, class and decision in the LCT.

In Fig. 4A, classes with the same initial stimulus overlapped, specifically c1 with c2 (P1 = G) and c3 with c4 (P1 = E), with c2 and c4 colors arbitrarily on top. P2 marginalization in Fig. S5 A revealed classes sharing the second stimulus merging, for example, c1 with c3 (P2 = G) and c2 with c4 (P2=E), with c3 and c4 colors arbitrarily plotted on top. Decision-based marginalization in Fig. 4B combined classes with matching outcomes, resulting in c1 with c4 and c2 with c3 overlapping, with blue and green traces arbitrarily on top. Class marginalization in Fig. 4C showed a similar merging (colors and class matches), specifically in the emergent sensory dynamics from class-dPC1.

### Uniform Manifold Approximation and Projection (UMAP)

This algorithm is a non-linear dimensionality reduction technique, based on: (1) approximating the manifold on which we assume the data was sampled from and (2) projecting this manifold into a low dimensional space while maintaining its local and global topological properties (28). For this, we concatenated all the 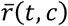 vectors horizontally for each neuron, according to the TPDT or LCT set. The firing rate was calculated using a window of 100 ms and steps of 50 ms, yielding a total of 170 time-bins per class when considering the whole TPDT or LCT (−1 to 7.5 s, Figs. S4 and S5) or 140 time-bins per class when considering a restricted TPDT or LCT (−1 to 6 s, without decision, Figs. 5 and 6), and 680 or 560 dimensions once all vectors were concatenated, when considering the whole or restricted task variants, respectively. As a result, the size of the matrix input into UMAP was 1646x680 or 1646x560 for the entire and restricted TPDT, respectively, and 313x680 or 313x560 for the entire and restricted LCT, respectively, where the 1646 (TPDT) or 313 (LCT) neurons were taken as observations, and the 680 or 560 time-point dimensions were reduced to two representative dimensions. We then project the inferred manifold into the new 2-dimensional space to determine whether a non-linear dimensionality reduction technique could isolate (or not) neural responses with the characteristic population dynamics identified in Figs. 3, S3 and 4. In Figs. 5B, 6B, S4A and S5B, we included a density contour plot calculated with a Gaussian kernel (σ = 0.5). For Figs. 5A, 6A, S4B and S5A we calculated a double-Gaussian (σ = 0.5) weighted-average centered on each of the colored X marks to visualize how the dynamics vary across the UMAP plane: neurons were weighted based on the Gaussian Probability Density Functions and then divided by the total sum of the weights.

### Network model

S2 neural population dynamics was modeled as nonlinear recurrent neural network (RNN)

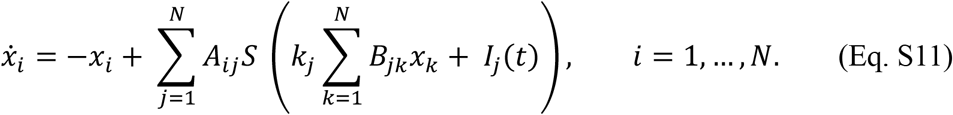

where *x*_*i*_(*t*) describes the average activity (firing rate) of a neuron *i* in a temporal window centered at time *t*. Depending on the context, *x*_*i*_(*t*) can also be interpreted as the average activity of a sub-population *i* of neurons. In vector form, model (1) becomes:

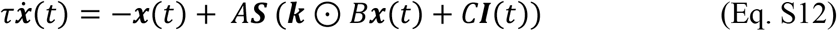

where ***x*** = (*x*_1_,…, *x*_*N*_), ***I*** = (*I*_1_,…, *I*_*N*_), ***k*** = (*k*_1_,…, *k*_*N*_), ***S***(***x***) = (tanh(*x*_1_),…, tanh(*x*_*n*_)), and ⊙ denotes the Hadamard product. The sigmoidal function *S*(*⋅*) = tanh(*⋅*) models neurotransmitter release and other kind of synaptic and intrinsic neural nonlinearities. Matrices *A* and *B* encodes the recurrent interconnection topology. *B*_*jk*_ is the effect that the activity of a neuron (or neural sub-population) *k* has on the liberation of neurotransmitter *j. A*_*ij*_ is the effect that the liberation of neurotransmitter *j* has on the activity of neuron (or neural sub-population) *i*. The gains *k*_*j*_ ≥ 0 control the overall strength of neurotransmitter release, thus controlling the linearized recurrent interconnection strength 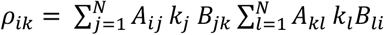 between neurons (or neural sub-populations) *i* and *k*. The eigenvector of the linearized recurrent interconnection matrix *A*(***k*** ⊙ *B*) are the network eigenmodes determining the qualitative network response stimuli.

Our network design procedure involves the following steps. 1. Fix the gains *k*_*j*_, which fixes the eigenmode types, as detailed in the main text. 2. Fix a family of (independent) eigenvectors *v*_*j*_, which fix the eigenmode directions in the space state. In particular, the *i*-th component of eigenvector *v*_*j*_ is the contribution of neuron (or neural sub-population) *i* to eigenmode *j*. 3. Define the change of variable matrix *U* = [*v*_1_ … *v*_*N*_]. 4. Let *A* = *U* and *B* = *U*^-1^ *in* (2), which enforces that the network model (2) has eigenmode *v*_*j*_ with eigenvalues *k*_*j*_.

Based on the step above, we generated a meta-population of neurons by running *n*_*rep*_ = 50 instances of an *N* = 6 neuron/neural sub-population network with the following parameters. Let 𝒰(*a, b*) denote the uniform distribution on the interval (*a, b*) and 𝒩(*μ, σ*) the Gaussian distribution with mean *μ* and variance *σ*. Active trial: *k*_1_, *k*_3_, *k*_4_ ∈ 𝒰(1.025, 1.05), *k, k*_5_ ∈ 𝒰(0.95, 1.0), *k*_6_ ∈ 𝒩(1.0, 0.0025). Hence, 6 is predominantly temporal. The eigenmode directions were generated according to the same distributions in both active and control trials as follows. Let *α*_1_, *α* ∈ 𝒰(0,1) and *β*_1_ = 1 − *α*_1_, *β* = 1 − *α*_2_. Then *v*_1_ = (α_1_, *β*_1_, 0, 0, 0, 0), *v* = (*β*_1_, *α*_1_, 0, 0, 0, 0), *v*_3_ = (0, 0, 1, 0, 0, 0), *v*_4_ = (*α*_2_, *β*_2_, 0, 0, 0, 0), *v*_5_ = (*β*_2_, *α*_2_, 0, 0, 0, 0), *v*_6_ = (0, 0, 0, 0, 0, 1), in such a way that there are 4 mixed eigenmodes (1, 2, 4, 5) and two pure eigenmodes (3, 6). Observe that the generated eigendirections are independent with probability one. In active trials, categorical eigenmodes were also receiving a constant inhibition with intensity *I*_*inh*_ = 0.0075 that favored meta-stability instead of bistability of the mode. Eigenmode 1, 3, 4 were receiving the stimuli with intensity *I*_*stim*_ = 0.0666, eigenmodes 2, 5 with intensity *I*_*stim*_ = 0.05, and eigenmode 6 with *I*_*stim*_ = 0.01. Julia code used to generate the numerical results, is available upon request.

## Data availability

Data files are publicly available at Zenodo (DOI: 10.5281/zenodo.4421855); see (51). Source data are provided with this paper.

## Code availability

The custom python and C scripts employed in the analysis of this data, as well as the experimental protocols, are available from the corresponding authors on reasonable request.

---

**Supplementary Figure 1.**
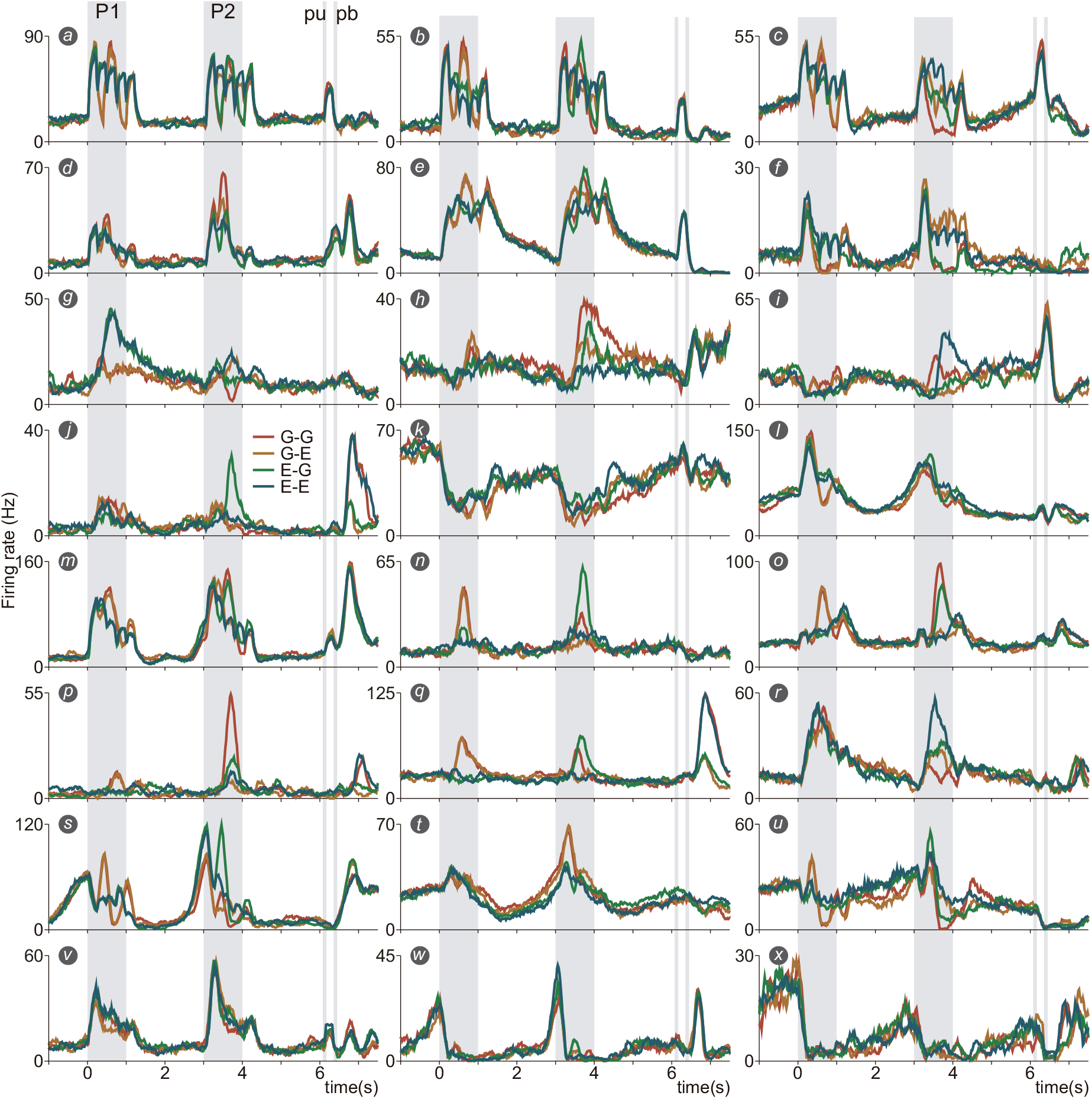
Activity of single neurons in S2. A mix of representative neurons (24 units in total, from *a* to *x*), wherein sensory, categorical, and temporal responses, are shown during TPDT. Traces represent firing rate averages per-class for each neuron. Each color refers to one of the four possible pattern pairs (class) of G (grouped) and E (extended): c1 (G-G, red), c2 (G-E, orange), c3 (E-G, green) and c4 (E-E, blue).

**Supplementary Figure 2.**
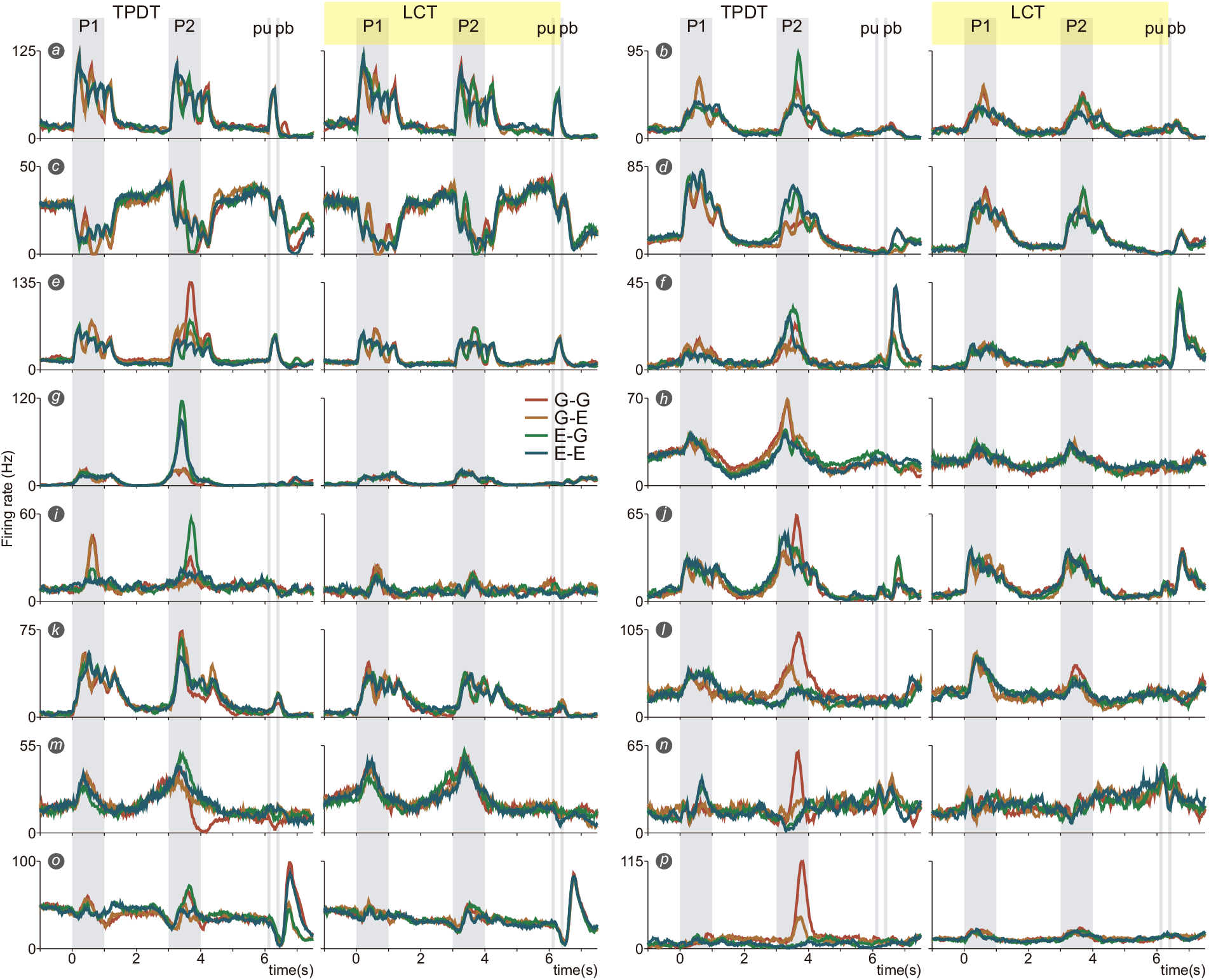
Single S2 neurons recorded during the TPDT and LCT. Exemplary neurons (16 units, from *a* to *p*, shown in TPDT and LCT) exhibited sensory, categorical, and temporal responses. Traces represent firing rate averages per-class for each neuron in both tasks. Each color refers to one of the four possible pattern pairs (class) of G (grouped) and E (extended): c1 (G-G, red), c2 (G-E, orange), c3 (E-G, green) and c4 (E-E, blue). Notice how neurons, as units *g and p*, turned off their categorical coding during the control task, leaving only temporal representations.

**Supplementary Figure 3.**
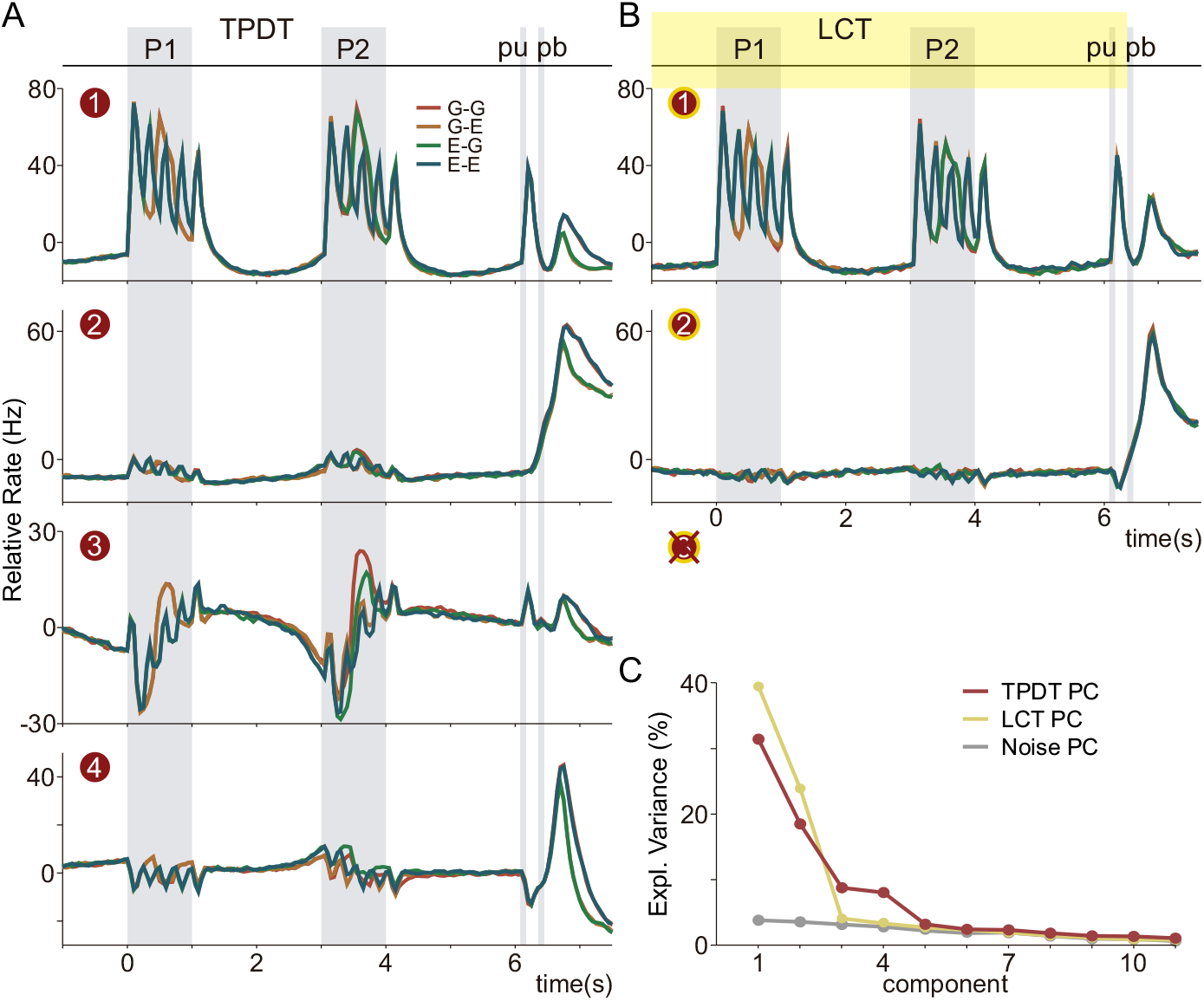
Population dynamics in S2 during the whole TPDT and LCT. (A, B) Principal component analysis (PCA) was applied to the covariance matrices computed with the neural activity from the entire active and control task (−1 to 7.5 s), to derive their principal components (PCs). Then the activity from the whole TPDT (n=1646) and LCT (n=313) sorted by class identity were projected onto those PCs and ordered by their explained variance (PC1: 31.44%; PC2: 18.5%; PC3: 8.75%; PC4: 8.02%), which are also resumed in panel C. In TPDT, sensory (PC1), transformed/categorical (PC3) and inter-trial decision (PC3 & PC4) dynamics plainly emerged. On the other hand, in LCT, only sensory (PC1) and temporal inter-trial (PC2) dynamics, remained.

**Supplementary Figure 4.**
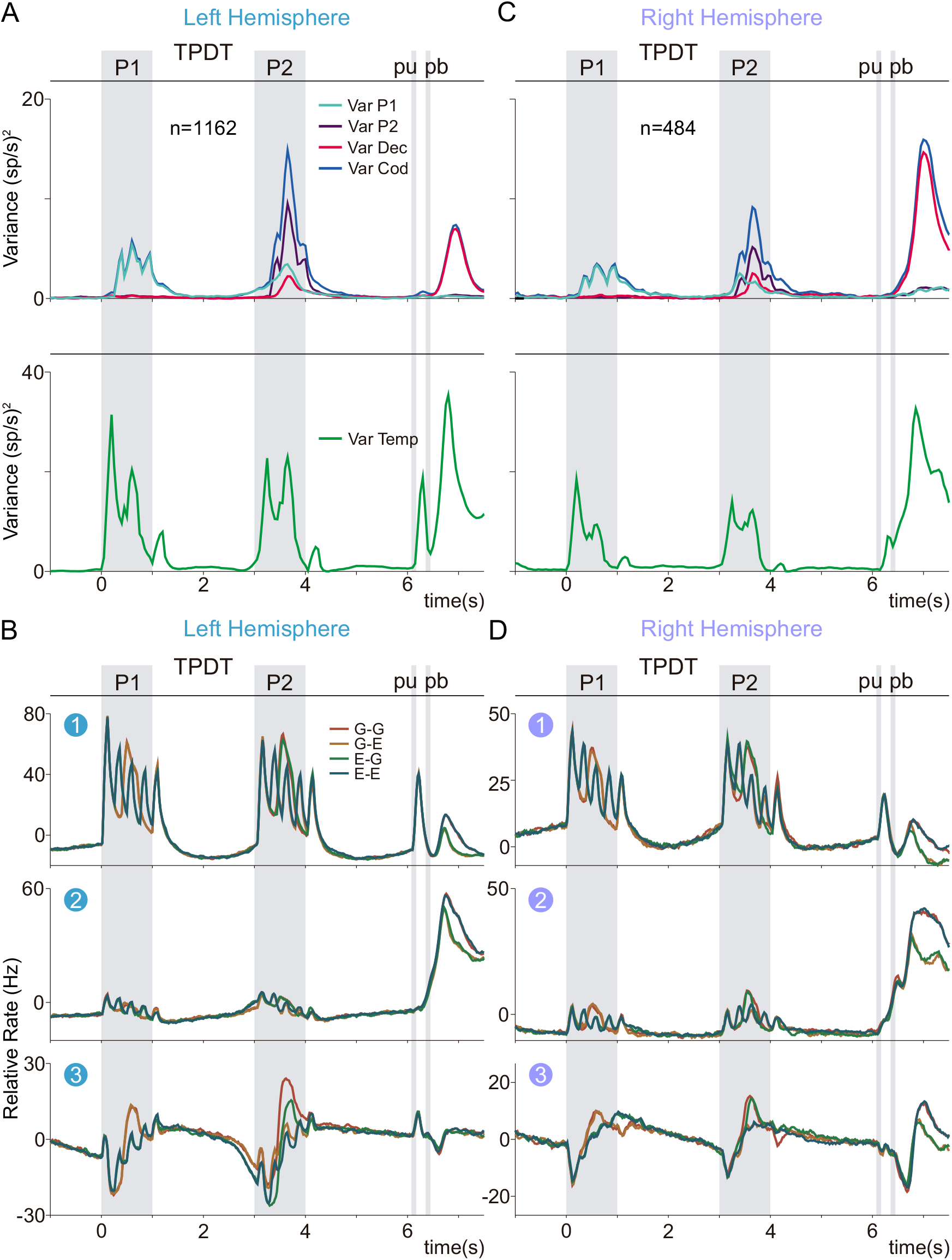
Cognitive variances and population dynamics across both hemispheres of the S2 Cortex. (A, C) Top: population variances as a function of time on the left (n=1162) and right (n=484) hemispheres, respectively. Traces refer to P1 (*Var P1*, light blue trace), P2 (*Var P2*, purple trace), decision (*Var Dec*, pink trace) and coding (*Var Cod*, blue trace) variances. All residual fluctuations were subtracted in all variances computed. Note the slight increase in the sensory component in the left hemisphere (ipsilateral to stimulation) and the marked increment in decision-related signals in the right one (ipsilateral to the motor execution). Bottom: temporal (or non-coding) variance (*Var Temp*, green trace) in the left and right hemispheres, respectively. This variance, which primarily captures the sequential events of the task, also showed a prominent sensory component during P1 and P2 in the left hemisphere and a more sustained activity during the decision period in the right one. (B, D) PCA was applied to the covariance matrices computed with the neural activity of the entire TPDT (−1 to 7.5 s) recorded in the left and right brain hemispheres, respectively, to derive their PCs. Then the activity from the whole TPDT in the left (n=1162) and right (n=484) brain hemispheres, sorted by class identity, were projected onto those PCs and ordered by their explained variance. Note how in both hemispheres sensory (PC1), categorical (PC3) and decision-related (PC3) dynamics plainly emerged, being more prominent the sensory component in the left one and the decisional in the right one.

**Supplementary Figure 5.**
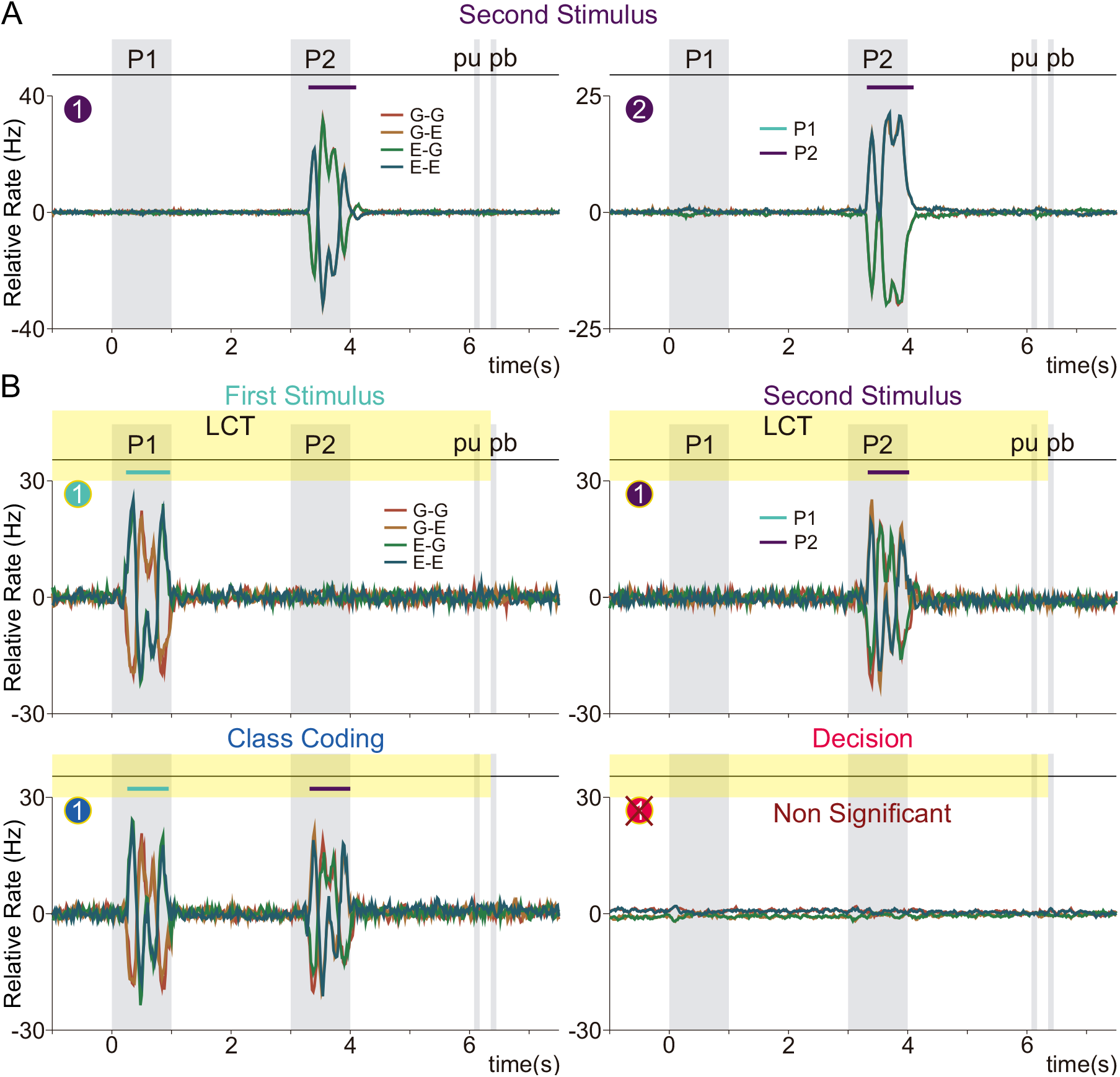
Demixed orthogonal dynamics in P2 and in LCT. (A) dPCA was applied to the marginalized covariance matrix from the whole TPDT (−1 to 7.5 s) with respect to P2 to derive the associated dPCs (P2-dPC1 & P2-dPC2). Population activity for TDPT (n=1646), sorted by class identity, was projected onto the resulting dPCs axes, and ordered by their EV. Only the first two dPCs for both parameters were above the noise level. Color lines on top of each projection represent significant coding periods. Projections exhibited purely sensory dynamics in the P2-dPC1 and categorical ones in the orthogonal component (P2-dPC2). (B) dPCA was applied to the marginalized covariance matrices from the whole LCT (−1 to 7.5 s) with respect to P1 (top left), P2 (top right), class coding (bottom left) and decision (bottom right) to derive their most significant dPCs. Only one dPC emerged above the noise level for P1, P2, and class coding, with none identified for decision coding. Population activity for LCT (n=313), sorted by class identity, was then projected onto those resulting dPCs axes. Color lines on top represent significant coding periods. Note that representations displayed in the only axes for P1, P2 and class coding yielded purely sensory dynamics.

**Supplementary Figure 6.**
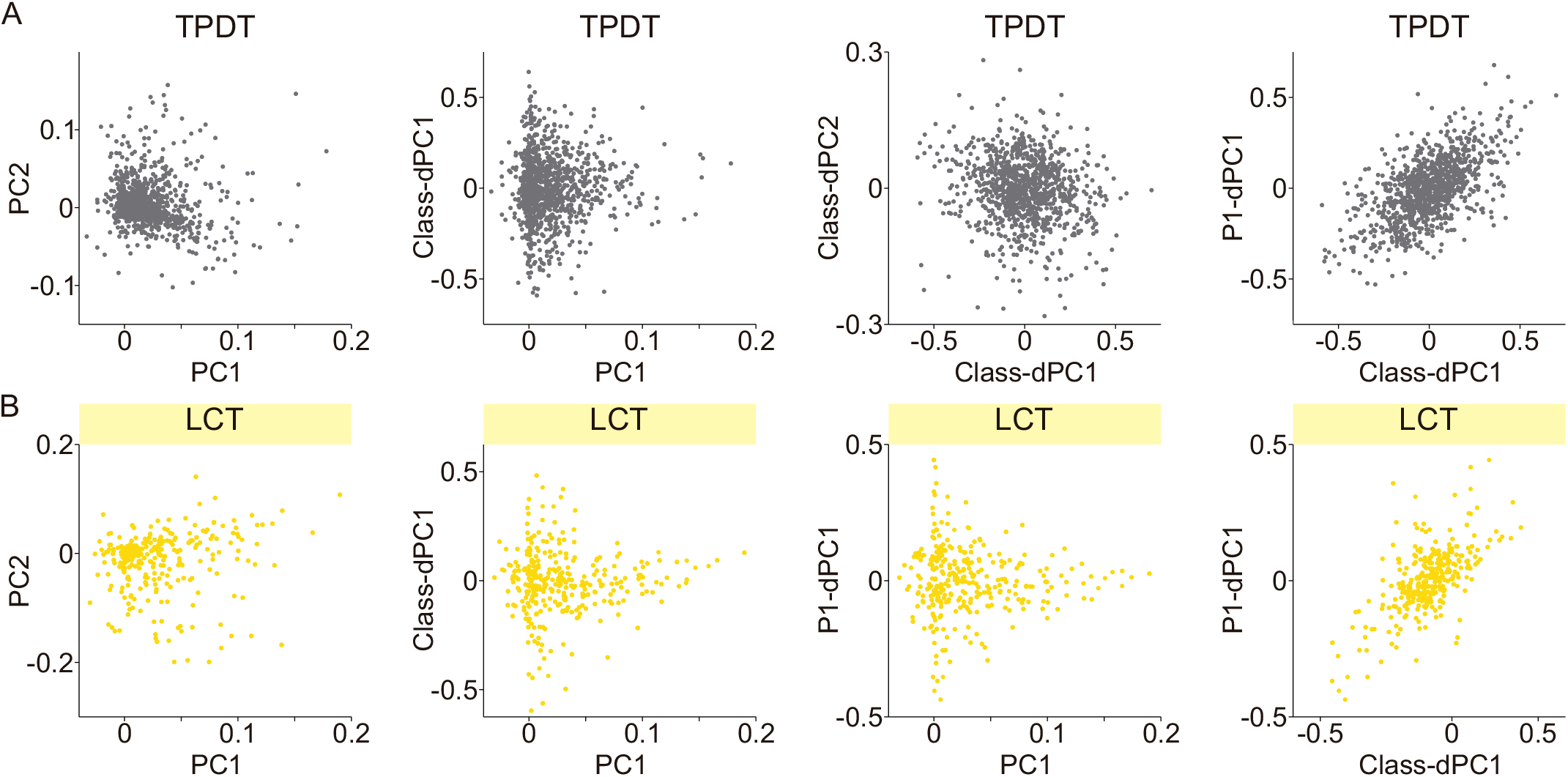
Continuity in the S2 network through linear approximations (PCA and dPCA). (A) Neuronal weights captured by the most prominent axes derived by applying PCA (PC1 & PC2) or dPCA (class-dPC1 & 2 and P1-dPC1) to the corresponding covariance matrices from the whole TPDT (−1 to 7.5 s) were plotted within each (PC1 vs PC2; class-DPC1 vs class-DPC2; class-dPC1 vs P1-dPC1) or across (PC1 vs class-dPC1) linear methods, to examine the existence of clusterization in the neural substrate. Neither of these plottings exhibited the presence of activity clusters across all S2 recorded neurons in this condition. (B) same procedure than in (A) but using the neuronal weights from the most significant axes derived by applying PCA (PC1 & PC2) or dPCA (class-dPC1 and P1-dPC1) to corresponding covariance matrices obtained from the whole LCT (−1 to 7.5 s). The plotting within (PC1 vs PC2; class-dPC1 vs P1-dPC1) or across (PC1 vs class-dPC1; PC1 vs P1-dPC1) linear methods, also revealed no evidence of cluster formation within S2 neural activity under this experimental condition.

**Supplementary Figure 7.**
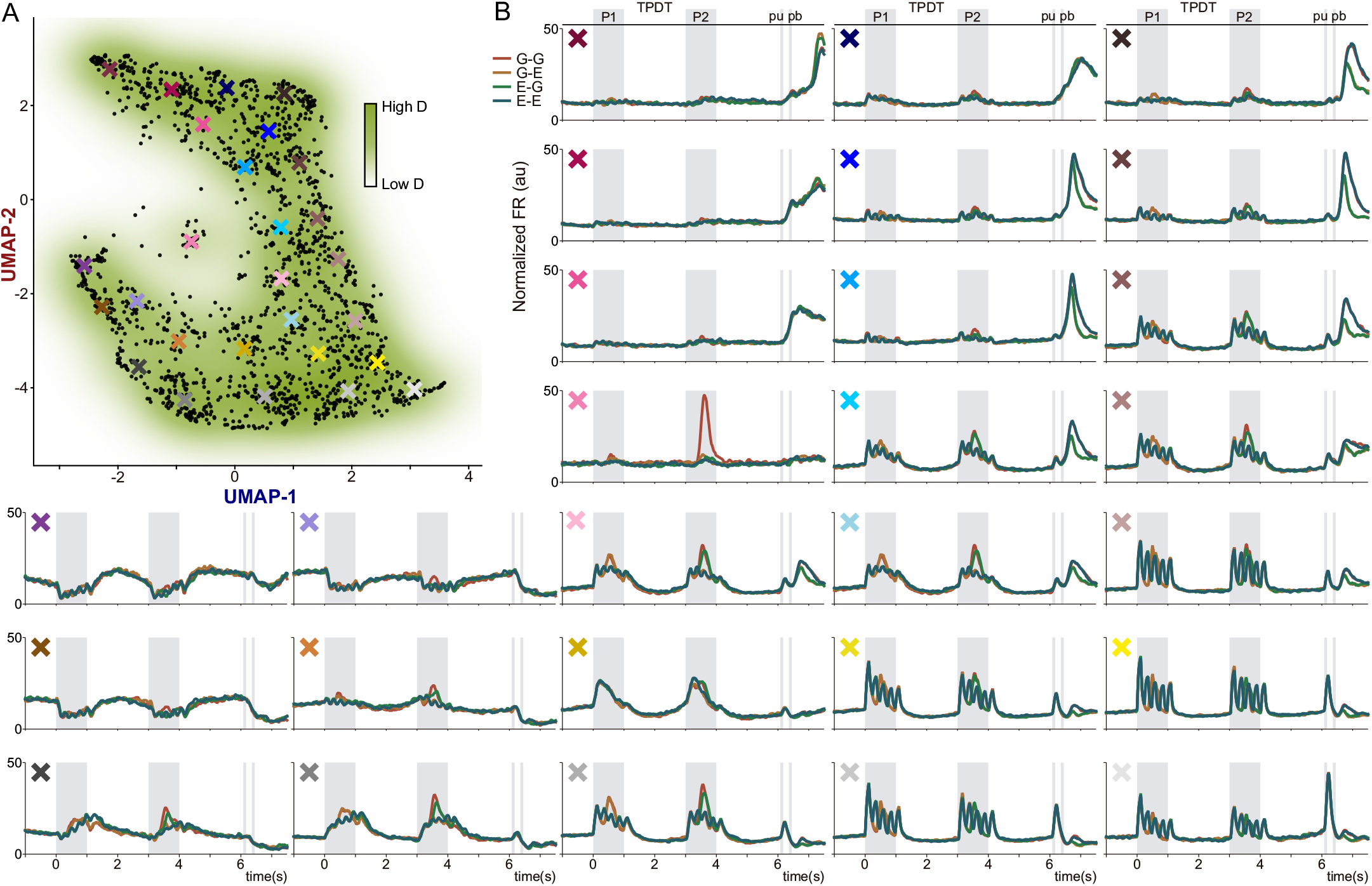
Orthogonal dynamics emerged in S2 from continuum responses during the TPDT. Contrary to Figure 5, here, we employed the neural activity from the whole TPDT. (A) 2-D nonlinear UMAP density projection of the normalized activity of all units (n=1646, each black dot) recorded during the whole TPDT (−1 to 7.5 s), exhibited a continuum of responses. The color X marks represents the centers of double-Gaussian (σ = 0.5) weighted averages displayed in the panels in B. (B) Each subpanel represents the weighted average of neuronal activity and is presented in back-transform arbitrary units that roughly correspond to the firing rate values (Hz). Notice that categorical average responses are surrounded by sensory and temporal average signals, as they also are anchored in the middle of the 2D projection. This kind of response might be useful for the S2 network to connect and form the continuum.

**Supplementary Figure 8.**
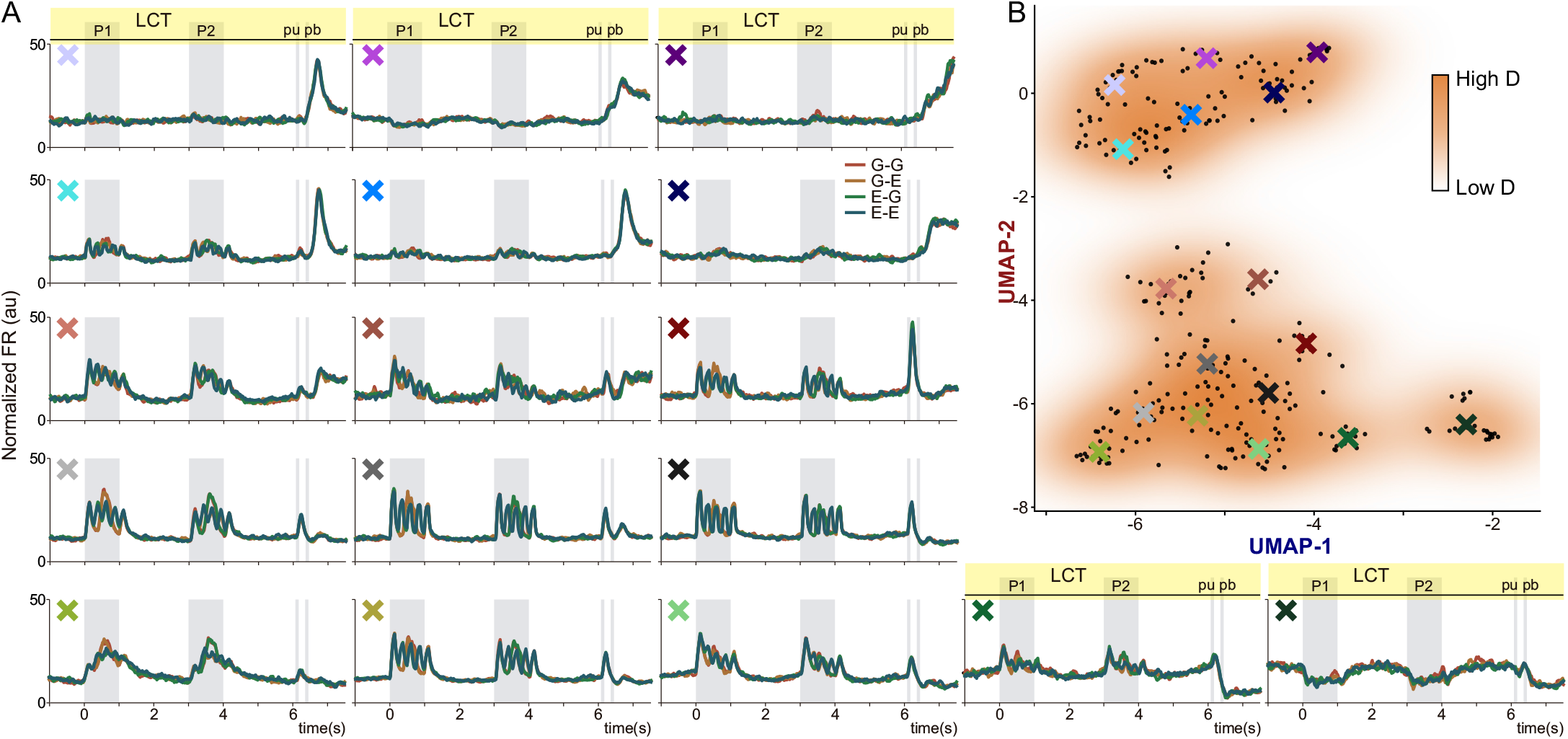
Continuum of responses in S2 breaks under the control task. Contrary to Figure 6, here, we employed the neural activity from the whole LCT. (A) Each subpanel represents the weighted average of neuronal activity in arbitrary units that approximately correspond to firing rate values (Hz). As LCT is a task variant that does not demand cognitive effort, the representative averages displayed only sensory and temporal responses. (B) UMAP density projection of the normalized activity of all units (n=313, each black dot) recorded during the whole LCT (−1 to 7.5 s), showed a breakup in the continuum that emerged during TPDT, exhibiting discrete subclusters of sensory (bottom) and temporal (top) dynamics. The color X marks represents the centers of double-Gaussian (σ = 0.5) weighted averages that are displayed in the panels in A.

**Supplementary Figure 9.**
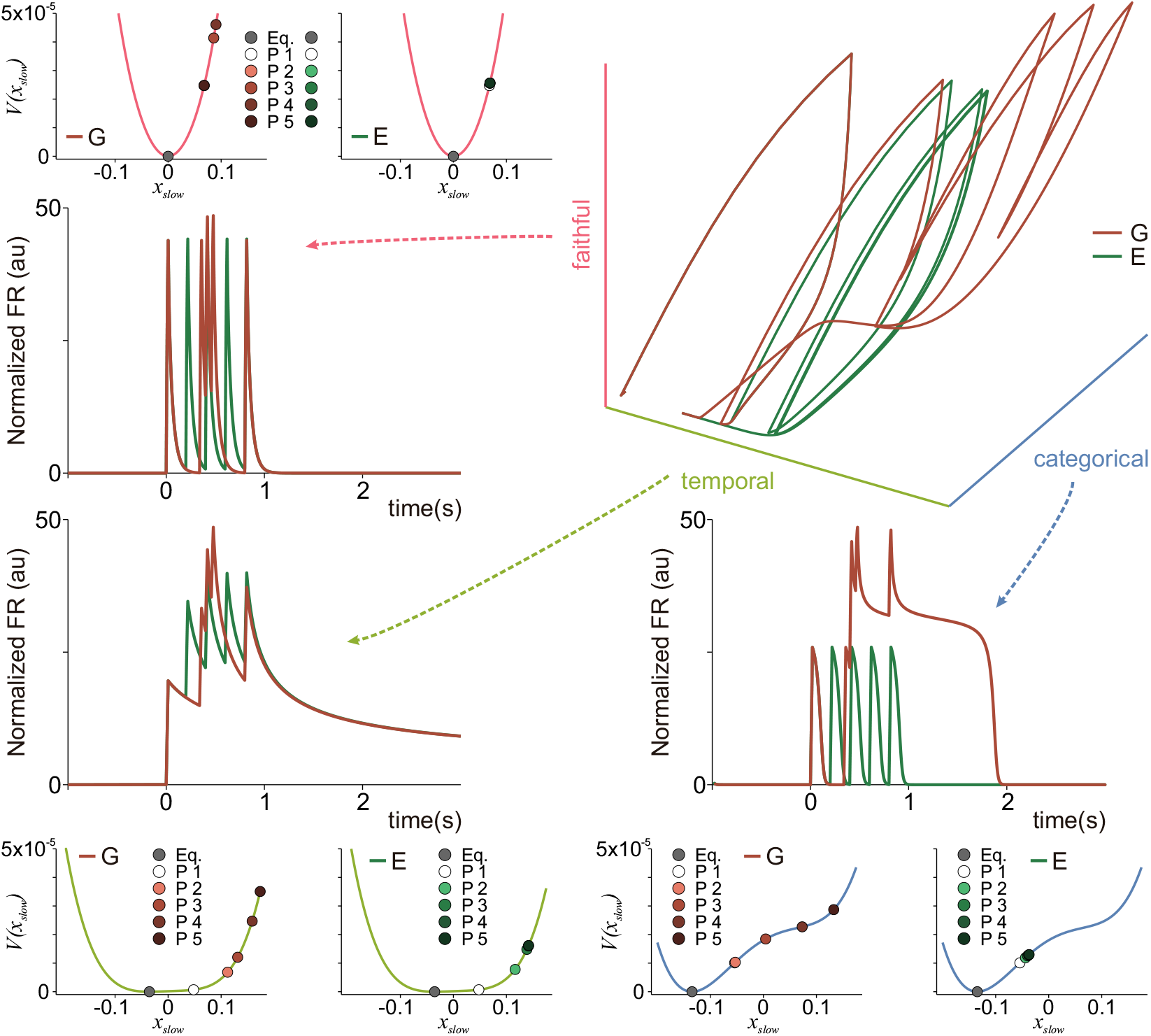
Eigenmodes define network dynamics in S2. Encoding along different network eigenmodes is determined by the qualitative properties of the dynamics inside each eigenmode. The three different eigenmode types of the simulated S2 network: faithful (pink), temporal (orange), categorical (dark blue). For each mode, we show the associated one-dimensional response and sketch the 1D potential function that determines the mode qualitative dynamical response. Faithful modes are characterized by a small network feedback gain and therefore their dynamics are dominated by a strongly stable response, associated with a sharply concave potential function, which makes the mode information encoding fast, strongly attractive, and faithfully tracking the presented stimulus. Temporal modes are characterized by an intermediate strength of network feedback, which creates a source of recurrent amplification, and therefore have much a flatter potential function, which slows down the response and allows for temporal integration of the presented stimuli. Categorical modes are characterized by a large network feedback gain and therefore are locally dominate by recurrent amplification, which leads to a non-convex potential function, with either a double well (bistable case) or not (metastable case), whose inflection point determines the appearance of a threshold for categorization.

